# Genetic landscape of DNMT3A R882H reveals mechanism of aberrant oligomerization

**DOI:** 10.1101/2025.09.24.678339

**Authors:** Emma M. Garcia, Olivia R. Lavidor, Shaunak Raval, Nicholas Z. Lue, Jessica K. Liang, Maeve McFadden, Derek D. Hwang, Marcanthony U. Zepeda, Luke W. Chen, Steven A. Carr, Malvina Papanastasiou, Brian B. Liau

## Abstract

DNA methyltransferase 3A (DNMT3A) is a *de novo* DNA methyltransferase that is recurrently mutated in hematological malignancies and developmental disorders. The most prevalent mutation, R882H, compromises DNMT3A activity in a dominant-negative manner, but its precise biochemical mechanism has been debated. Here, we use paired deep mutational scanning of the wild-type and R882H-mutant proteins to systematically identify mutations on a massively parallel scale that modify DNMT3A activity by suppressing, phenocopying, or selectively rescuing the dominant-negative effect of R882H. By leveraging the mutational depth and unbiased nature of the paired genetic landscapes, we uncover two distinct mechanisms that can rescue DNMT3A^R882H^ activity, providing novel insights into the function of the R882 hotspot. First, by analyzing the effects of combinatorial mutations in the target recognition domain (TRD), we reveal that its crosstalk with the ADD regulatory domain modulates DNMT3A DNA binding and enzymatic activity, partially compensating for R882H-induced loss-of-function. Second, pairwise analysis of variant effects across the two genetic backgrounds supports the notion that R882H promotes aberrant macro-oligomerization of DNMT3A via its central dimerization interface, which accounts for its dominant-negative effect. Critically, we show that the R882 position exhibits a distinct dominant-negative signature in the genetic landscape, where positively charged residues at this position safeguard against aberrant macro-oligomerization. By performing hydrogen-deuterium exchange mass spectrometry (HDX-MS) on DNMT3A mutants, we show that R882H dramatically alters the protein dynamics of DNMT3A, rigidifying its central dimerization interface to promote oligomerization. Our data support a new model in which R882H removes the critical function of R882, where the arginine attenuates the pre-organization of the interface and subsequent oligomerization at a supramolecular assembly hotspot^1^. Altogether, we map the genetic landscape underpinning the DNMT3A^R882H^ hotspot mutation, illuminating the unexpected molecular mechanism of macro-oligomerization that drives its dominant-negative effect.

## Introduction

DNMT3A is an essential de novo DNA cytosine methyltransferase critical for gene regulation and hematopoiesis. The R882 position of *DNMT3A* is the most common somatically mutated locus in human blood^2,3^. In acute myeloid leukemia (AML), more than 20% of patients harbor a mutation in *DNMT3A*, ∼65% of which involve R882^4^. R882 mutations are typically heterozygous and the most frequently observed mutation, R882H, has been shown to exert a dominant-negative effect on the methyltransferase activity of DNMT3A^5,6^. Loss of DNMT3A reprograms DNA methylation and promotes the aberrant clonal expansion of hematopoietic stem and progenitor cells ^7^. Because it is dominant negative, the R882H mutation compromises DNMT3A activity more than other single-copy, loss-of-function mutations, leading to its higher frequency in patients ^7^. Due to its clear biological and clinical consequences, the molecular mechanism of DNMT3A R882H has been the focus of intense study^5–18^.

DNMT3A and its paralog DNMT3B both contain three protein domains: a catalytic methyltransferase (MTase) domain and the regulatory ADD and PWWP domains, which recognize histone marks to target DNA methylation to specific genomic regions (**Figure 1A**). Both DNMT3A and DNMT3B methylate cytosines as active homo- or hetero-tetramers^19,20^ and multimerize using two distinct interfaces: (1) the hydrophobic FF interface, and (2) the central RD interface, which also binds DNA^8,9,19,21^. While the FF interface is largely dominated by hydrophobic contacts, the RD interface is predominantly mediated by electrostatic interactions, including the eponymous R885–D876 salt bridge. Many mutations associated with clonal hematopoiesis and AML, including the hotspot R882H, are found at the RD-RD protein-protein interface^4,22^, highlighting its functional importance.

**Figure 1:**
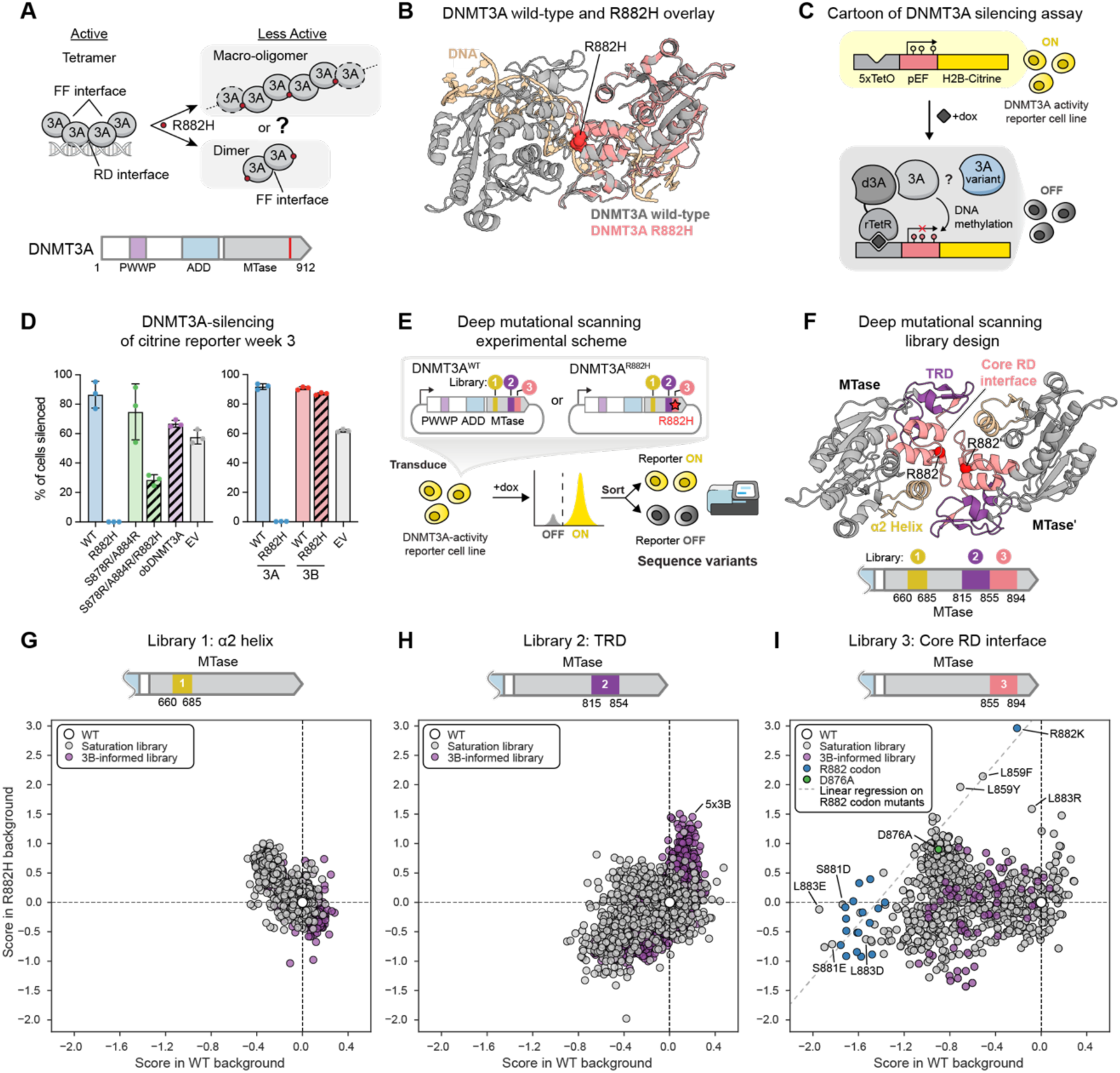
Paired deep mutational scanning of DNMT3A RD interface. **A.** Top: Schematic of DNMT3A oligomer formation by the FF and RD interfaces and competing models for the dominant negative effect of R882H. Bottom: Domain map of DNMT3A. MTase = methyltransferase. **B.** Structural overlay of two, RD interface-forming copies of wild-type (grey) and R882H (salmon) DNMT3A MTase domains bound to DNA (gold). Only one copy of DNMT3A^R882H^ is depicted for clarity. PDB: 6w89, 6w8b. **C.** Schematic of citrine reporter assay to measure DNMT3A activity in live cells. 3A = endogenous DNMT3A, 3A variant=exogenously overexpressed DNMT3A copy, d3A = DNMT3A^E756A^ (catalytically inactive DNMT3A). **D.** Flow cytometry quantification of silencing (percent dark) for reporter cells expressing indicated DNMT3A variants, DNMT3B variants, or empty vector with 1 mg/mL dox treatment after 21 days. Data are mean ± S.D. with individual replicates shown (n=3). **E.** Schematic of paired deep mutational scanning on DNMT3A^WT^ and DNMT3A^R882H^. Reporter cells expressing a surface marker and citrine are individually transduced with one of six libraries (libraries 1-3, with or without R882H) treated with dox, then sorted for reporter silencing and sequenced. For a more detailed schematic see **Figures S3C and S3D**. **F.** Structural view showing regions of the RD interface mutated in the deep mutational scan. TRD = target recognition domain. PDB: 6w8b. **G.** Scatter plot of deep mutational scanning scores for DNMT3A^WT^ (*x* axis) or DNMT3A^R882H^ (*y* axis) for Library 1 (a2 helix) normalized to wild-type or R882H, respectively. Data are mean of n=3 replicates. Grey = variants from saturation library; purple = variants from DNMT3B paralog-informed library. **H.** Scatter plot of deep mutational scanning scores for DNMT3A^WT^ (*x* axis) or DNMT3A^R882H^ (*y* axis) for Library 2 (TRD). Data are mean of n=2 replicates (WT) or n=3 replicates (R882H). **I.** Scatter plot of deep mutational scanning scores for DNMT3A^WT^ (*x* axis) or DNMT3A^R882H^ (*y* axis) Library 3 (Core RD interface). Data are mean of n=3 replicates. Blue = mutations at the R882 position, green = D876A. Grey dashed line indicates the linear regression of mutations at the R882 codon that were identical in the WT and R882H background libraries (R882H score = 2.279(WT score) + 3.284). Source data can be found in **Table S1.**

Despite its initial discovery over 15 years ago^23^, the precise biochemical mechanism by which R882H exerts its effect on DNMT3A has been debated. Two contrasting mechanistic models for the dominant-negative effect of DNMT3A^R882H^ have been proposed: that the R882H mutation (1) disrupts the central RD-RD interface to form dimers only mediated by the FF interface^5,10^ or (2) remodels the RD interface region to promote macro-oligomerization beyond tetramers, in either case trapping DNMT3A in an inactive, non-tetrameric form (**Figure 1A**)^8,11^. Additionally, R882H reduces DNMT3A’s binding to DNA and overall enzymatic activity and perturbs its flanking sequence preference^9,11–13,16,17^. Consistent with an oligomerization model, recent high-resolution structures of the MTase domains of DNMT3A^WT^ and DNMT3A^R882H^ suggest that R882H-specific protein-protein contacts involving the histidine glue together the central RD interface. However, these and other crystal structures^8,9^ revealed only minor differences between the wild-type and mutant RD protein-protein interface (**Figure 1B**). Critically, these landmark structures of DNMT3A could only resolve the interface residues when bound, strongly suggesting that the mechanism for R882H-driven oligomerization is rooted in the dynamics of the unbound state, which have remained elusive to crystallographic analysis.

Determining the molecular mechanism by which the R882H hotspot mutation exerts its dominant-negative effect would deepen our understanding of blood malignancies and, more broadly, how protein-protein interactions are corrupted in disease, informing future therapeutic strategies. In this pursuit, we used deep mutational scanning of DNMT3A^WT^ and DNMT3A^R882H^ to systematically chart the genetic landscape surrounding the R882H mutation on a massively parallel scale, profiling the activity of over 8,000 variants. By leveraging mutations that modify or phenocopy the dominant-negative effects of R882H on DNMT3A activity, our study supports the macro-oligomerization model. Comprehensive mutagenesis of R882 and its surrounding residues demonstrates that the positive charge of R882 safeguards against macro-oligomerization through negative design — a widely observed phenomenon but not yet linked to cancer mutation etiology^24^. Thus rather than R882H directly enhancing interactions at the RD-RD interface, our results instead indicate that the cancer mutation primarily drives DNMT3A oligomerization through loss of a critical safeguarding residue. In addition, unbiased mutational scanning identified unexpected and rare mutations that selectively rescue the effects of R882H. By examining these mutants in mechanistic studies, including HDX-MS, we show that the R882H mutation promotes aberrant macro-oligomerization of DNMT3A by rigidifying the RD interface, rendering it optimal for multimerization. Altogether, by employing the unbiased high-content data afforded by deep mutational scanning, we revise the mechanistic model of R882H-promoted oligomerization and uncover a novel mechanism for corrupting protein-protein interactions in human cancers.

## Results

### Paired deep mutational scanning reveals the genetic landscape surrounding DNMT3A R882H

We posited that high-throughput genetic approaches could provide insight into the dominant-negative mechanism of R882H by comprehensively identifying sequence variants that modify the activities of DNMT3A^WT^ and/or DNMT3A^R882H^. Towards this end, we first implemented a cell-based reporter assay that can measure the activity of DNMT3A and faithfully recapitulate the dominant-negative effect of the R882H allele. To do so, we adapted a doxycycline (dox)-inducible DNMT3A system we previously established^14,25,26^ in which a catalytically dead copy of DNMT3A (d3A) recruits endogenous DNMT3A^WT^ (3A) and/or exogenously expressed DNMT3A^variant^ (3A variant) to a reporter construct that drives citrine expression (**Figure 1C**). If DNMT3A recruitment is successful and the enzyme is active, it methylates the promoter and silences the citrine reporter. Due to the presence of endogenous DNMT3A in the reporter cells, a baseline level of ∼60% cell silencing is observed after 21 days of dox treatment (**Figure 1D**). As anticipated, additional overexpression of DNMT3A^WT^ accelerates citrine silencing. By contrast, overexpression of DNMT3A^R882H^ abrogates silencing, consistent with the mutant protein inactivating endogenous DNMT3A via a dominant-negative mechanism.

We next evaluated DNMT3A homologs using this cell-based reporter, reasoning that this might provide insight into the mechanism of R882H. First, we considered naturally occurring DNMT3A variants and generated a phylogenetic tree of *DNMT3* homologs containing a PWWP, ADD, and MTase domain (**Figure S1A**). R882 is highly conserved across both DNMT3A and DNMT3B branches of the tree but is not conserved in DNMT3 in invertebrates (**Figures S1B, and S1C**). Within this group, the residue equivalent to H882 is found in very few sequences, including the *DNMT3* sequence of *octopus bimaculoides* (*obDNMT3*). We reasoned that if obDNMT3 is a functional methyltransferase, its additional sequence variation might compensate for the effect of R882H. The most chemically dissimilar residues between human DNMT3A (hsDNMT3A) and obDNMT3 nearby the R882 position are S878 and A884 (numbered by hsDNMT3A), both of which are arginine in obDNMT3 (**Figures S1D and S2A**). Both obDNMT3 and DNMT3A^R882H/S878R/A884R^ displayed higher activity in our fluorescent reporter assay than DNMT3A^R882H^ (**Figures 1D, S2B and S2C**). The same trend was observed in a similar fluorescent reporter cell line modified by knockout of endogenous DNMT3A (**Figure S2D**), demonstrating that these DNMT3A^R882H^ variants exhibit activity even in absence of endogenous DNMT3A. Recent studies demonstrated that DNMT3B activity ^27^ and multimerization ^8^ are not impacted by R823H, the equivalent mutation of R882H in this paralog. In agreement, we tested DNMT3B variants in our reporter and observed that DNMT3B^R823H^ is nearly as active as DNMT3B^WT^ and DNMT3A^WT^ (**Figures 1D, S2E,F**). Altogether, these results indicate that the loss of DNMT3A^R882H^ activity can be partially rescued by sequence variation derived from DNMT3A orthologs and paralogs, and that this effect can be measured by our reporter assay.

Motivated by the rescue afforded by the ortholog- and paralog-informed mutants, we leveraged our reporter system to systematically identify mutations that modify or phenocopy the dominant-negative effect of R882H. We performed paired deep mutational scanning of both DNMT3A^WT^ and DNMT3A^R882H^ (**Figure 1E**), in which the pair of mutational scanning libraries only differ by the absence or presence of R882H. We modified our reporter assay to include a cell surface marker in addition to citrine to allow higher throughput cell sorting with magnetic beads and/or FACS (**Figure S3A, see Methods**)^28^. To enable direct short-read Illumina sequencing, we split the pooled mutant library into three sub-libraries spanning unique regions of the DNMT3A RD interface. These sub-libraries covered: (1) the α2 helix and surrounding residues, comprising part of the S-adenosyl-methionine binding pocket and residues forming contacts across the RD interface (residues 660-685, tan); (2) the first part of the target recognition domain (TRD), including the TRD loop (residues 815-854, purple); and (3) the remaining TRD and central RD interface (residues 855-894, pink) (**Figure 1F**). Within their respective regions, each sub-library covered both saturation mutagenesis and combinatorial variants informed by the DNMT3B sequence (**see Methods** and **Figure S3B**), based on the observation that DNMT3B tolerates the R882H mutation and that DNMT3B paralog-inspired single and double mutations can partially rescue DNMT3A^R882H^ activity^8^ . Each sub-library was introduced separately into full-length sequences encoding DNMT3A^WT^ and DNMT3A^R882H^, creating paired libraries only differing at position 882, for a total of 4,020 single amino acid substitutions and 4,736 combinatorial variants across 105 residues of DNMT3A (**Figures 1E and S3C**).

We introduced the paired libraries of DNMT3A variants into our reporter cells, initiated DNMT3A variant recruitment with dox treatment, then sorted for cells that had silenced the citrine/surface marker reporter gene (**Figures 1E and S3D**). As expected, cells overexpressing DNMT3A^R882H^-based libraries silenced the reporter more slowly than cells expressing DNMT3A^WT^-based libraries (**Figure S3E**). Finally, we sequenced genomic DNA isolated from the sorted cells and calculated the wild-type adjusted log-ratio enrichment of each sequence in the silenced versus the unsorted population^29^ (from here on referred to as “enrichment scores”, **see Methods**). Scores correlated well across two (Library 2 wild-type background) or three (all other libraries) replicates (R^2^ = 0.47-0.92) (**Figure S4**). By comparing the effects of mutations in the wild-type versus R882H-based libraries, these data together provide a genetic landscape surrounding the R882H hotspot mutation (**Figures 1G–I**).

Comparing the score of each mutant in the DNMT3A^WT^ versus DNMT3A^R882H^-based libraries showed different trends in each sub-library, suggesting diverse mechanisms occurring across these protein regions. The most striking perturbations to DNMT3A were observed for mutations in the central RD interface (Library 3), which constitutes the majority of the protein-protein interface and includes the R882 position. More modest scores were observed for the screens on the TRD (Library 2) and α2 helix (Library 1), consistent with their locations more peripheral to the interface. We investigated the mutations spanning these protein regions to determine their respective mechanisms for modifying the action of R882H.

### Modulation of TRD-ADD contacts allows for increased DNA binding and partially rescues the activity of DNMT3A R882H

We first examined mutations in our DNMT3B paralog-informed libraries. Of the three DNMT3B informed libraries, only Library 2 (the TRD library) contained variants that significantly modified the activity of DNMT3A^R882H^ (defined as variant score > 2 S.D. above the mean of the library). Despite the overall modest scores observed in Library 1 (α2 helix library), one of the top mutants in this library was the previously reported DNMT3A^R676K/R882H^ variant. However, since DNMT3A^R676K/R882H^ did not display substantial activity in an individual silencing assay using our system (**Figure S5A**), we focused our analysis on variants in the TRD library.

All variants within the TRD, a critical region of the enzyme responsible for DNA recognition and binding, were modestly correlated between the DNMT3A^WT^ and DNMT3A^R882H^ deep mutational scans (Spearman π = 0.610) (**Figures 1H and S5B**). Notably, this correlation is predominantly driven by the DNMT3B paralog-informed variants, including those with positive scores in both the DNMT3A^WT^ and DNMT3A^R882H^-based libraries. Examining the relationship between enrichment score and number of mutations per variant in the DNMT3A^R882H^-based library, we observed that a combination of 5 of the 11 mutations tested (S828K, R836K, D845N, H847L, F851V) yields a near-maximum score (**Figure 2A**); we hereon refer to this set of five mutations as 5×3B. Additional mutations such as E817D, H821Y, and E854G were largely neutral in the R882H background, while the remaining three mutations (G822N, F827L, R831Q) were depleted from top ranked variants (**Figure S5C**). As single variants, the DNMT3B paralog-inspired mutants did not strikingly differ from other variants in the saturation library (**Figure S5D**). Across all mutant combinations, S828K, R836K, and D845N generally contributed positive overall effects on variant score (i.e., mutational effect score), while F827L and E854G tended to have negative effects (**Figure 2B**). However, all mutations showed a wide range of mutational effect scores, indicating that epistatic relationships likely exist across the 11 mutations tested.

**Figure 2:**
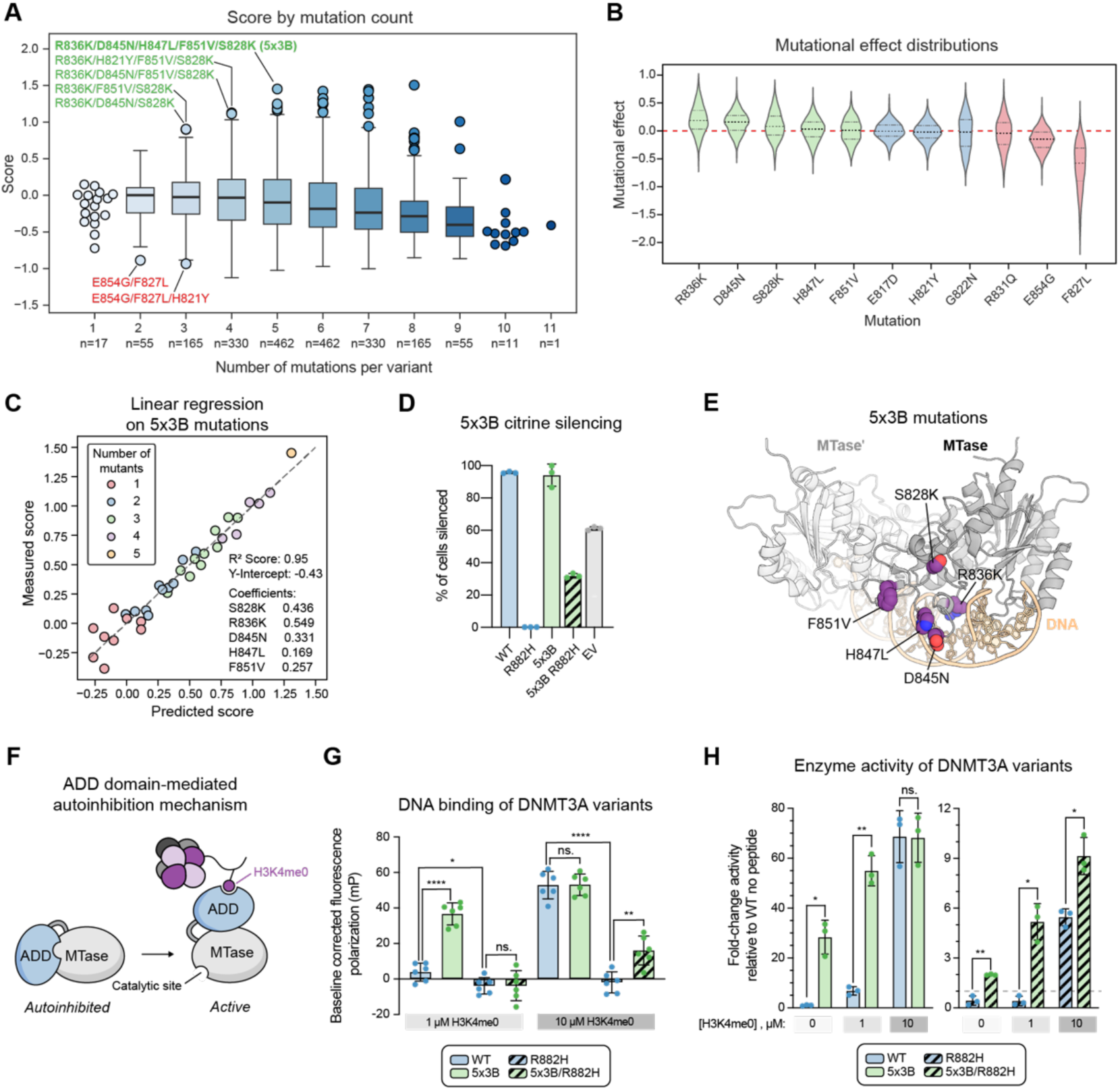
TRD mutations from the DNMT3B-informed library combine to relieve ADD autoinhibition and boost activity. **A.** Box plot of TRD DNMT3B-informed variant scores by number of mutations per variant. Select mutations of interest are labeled. center line, median; box, interquartile range; whiskers, up to 1.5 × interquartile range. **B.** Violin plot showing distribution of mutational effect scores for each of the DNMT3B informed mutations. Mutational effect is calculated as the difference in score between matched variants with and without the named mutation (e.g. DNMT3A^variant X^ versus DNMT3A^variant X/R836K^). Median and interquartile range are shown. Green = mutations in 5×3B, red = deleterious mutations, blue = other mutations. Mutations are numbered by DNMT3A residue. **C.** Linear regression analysis on variants containing the five mutations composing DNMT3A^5x3B^. Coefficients correspond to the predicted impact of each mutation on variant score. **D.** Flow cytometry quantification of reporter cells expressing indicated DNMT3A variants or empty vector (EV) with dox treatment after 21 days. Data are mean ± S.D. with individual replicates shown (n=3). **E.** Structural view of the MTase dimer (gray) bound to DNA (tan) with the mutated positions constituting 5×3B shown in spheres (purple = C, blue = N, red = O). PDB: 6w8b. **F.** Cartoon representation of ADD domain autoinhibition mechanism^33^. **G.** Fluorescence polarization of Cy3 labeled DNA with 1 μM of indicated DNMT3A variants in the presence of 1 or 10 μM H3K4me0. (H3K4me0-ADD K_D_ = 1.75 ± 0.11 µM)^33^. Experiment was performed in triplicate twice for a total of n=6. Data are mean ± S.D. with individual replicates shown. *P* values were calculated through two-tailed unpaired *t* tests; * ≤ 0.05, ** ≤ 0.01, *** ≤ 0.001, *** ≤ 0.0001. **H.** Quantification of the DNA methylation activity of indicated DNMT3A on a poly dI•dC substrate in the presence 0, 1, or 10 μM H3K4me0. Data are mean ± S.D. with individual replicates shown (n=3) and are representative of two independent experiments. *P* values were calculated through two-tailed Welch’s *t* tests; * ≤ 0.05, ** ≤ 0.01, *** ≤ 0.001, *** ≤ 0.0001. Data in **A**–**C** are mean of n = 3 biological replicates and the overall DMS experiment was performed once

In contrast to the epistatic behavior among all 11 mutations, linear regression on variants containing any subset of the 5×3B mutations showed consistent positive contributions to variant score for each mutation across the included backgrounds (**Figure 2C**), indicating that the effects of these mutations in this area of mutational space are additive. Notably, R836K showed the largest positive mutational effect score, both globally and in the linear regression of 5×3B variants. R836K has been previously implicated in specifying the sequence preferences, but not the activity, of DNMT3A versus DNMT3B^30–32^. Interestingly, the top-scoring mutant in the DNMT3A^WT^-based library is S828K/D845N/E854G while R836K appeared neutral (**Figures S5C and S5E**), indicating differing effects between the paired libraries. Individual validation in our cellular reporter assay showed that 5×3B increases the activity of both DNMT3A^WT^ and DNMT3A^R882H^, including in the absence of endogenous DNMT3A, consistent with the results of the screen (**Figures 2D, S5F, and S5G**).

We next investigated how 5×3B increases the activity of DNMT3A. Notably, prior studies have demonstrated that R882H impairs DNA binding and resulting enzymatic activity^9,11,13^. The most potent of the five single mutants, R836K, and two other mutations comprising 5×3B, D845N and H847L, are proximal to DNA as well as the interaction surface between the ADD and MTase domains (**Figure 2E**). Prior studies have shown that the ADD domain inhibits enzyme activity by blocking the association of DNA to the MTase domain and is released to an active conformation by binding to histone H3K4me0^33^ (**Figure 2F**). Consequently, we hypothesized that these mutations may alter DNA binding and/or the ADD-autoinhibition mechanism.

To test this hypothesis, we assessed the binding of purified DNMT3A variant proteins to fluorophore-labeled DNA in the presence of varying concentrations of H3K4me0 peptide using fluorescence polarization. As expected, R882H decreased DNA binding relative to the wild-type enzyme at both peptide concentrations (**Figures 2G and S5H**). DNMT3A^5x3B^ showed increased DNA binding relative to DNMT3A^WT^ in the presence of 1 μM peptide but similar levels with 10 μM peptide, suggesting that the mutations are disrupting autoinhibition but not the overall DNA binding affinity of the MTase domain. For DNMT3A^5x3B/R882H^, no binding was observed with 1 μM peptide and only low levels were observed with 10 μM peptide, consistent with 5×3B partially rescuing DNA binding in the R882H context. In agreement with these DNA binding trends, the 5×3B mutations also increased methyltransferase activity for both DNMT3A^WT^ and DNMT3A^R882H^ across all peptide concentrations tested except for DNMT3A^5x3B^ with 10 μM H3K4me0 (**Figure 2H**). Altogether, these results indicate that the 5×3B mutations partially relieve ADD-mediated autoinhibition in both the wild-type and R882H contexts to increase DNA binding, although we cannot rule out other mechanisms. Notably, neither the silencing nor DNA binding activity of DNMT3A^5x3B/R882H^ reached comparable levels to that of DNMT3A^WT^ (**Figures 2D, 2H, S5F, and S5G**), indicating that the activity of R882H could not be fully recovered by enhancing DNA binding.

### RD interface disruption suppresses the dominant-negative effect of R882H

We next investigated the deep mutational scans of the α2 helix (Library 1) and central RD interface (Library 3). Notably, in contrast to mutations in the TRD (Library 2), the effect of those targeting the α2 helix exhibited anti-correlation between the DNMT3A^WT^ and DNMT3A^R882H^ systems (Spearman ρ = –0.657). This anticorrelation was in part driven by a group of mutants that decreased DNMT3A^WT^ reporter silencing while increasing DNMT3A^R882H^ reporter silencing (**Figure 1G**). Variants with this behavior were also prevalent in the central RD interface library (Library 3) (**Figure 1I**). Many of these variants include nonsense mutations (i.e, premature stop codons) (**Figure 3A** (red outline) and **S6A**) which, as expected, decreased reporter silencing when introduced into DNMT3A^WT^. By contrast, nonsense mutations introduced in cis with R882H increased reporter silencing, consistent with removing the dominant-negative effect of DNMT3A^R882H^ and permitting endogenous DNMT3A to silence the reporter. Furthermore, mutations associated with protein destabilization, either as predicted by ThermoMPNN^34^ (a neural network predictor of protein stability) (**Table S2**) or demonstrated in previous studies^35^, exhibited similar effects to nonsense mutations (**Figures 3B and S6B–D**). Altogether, our results show that mutations that deplete DNMT3A protein levels suppress the dominant-negative effect of R882H.

**Figure 3:**
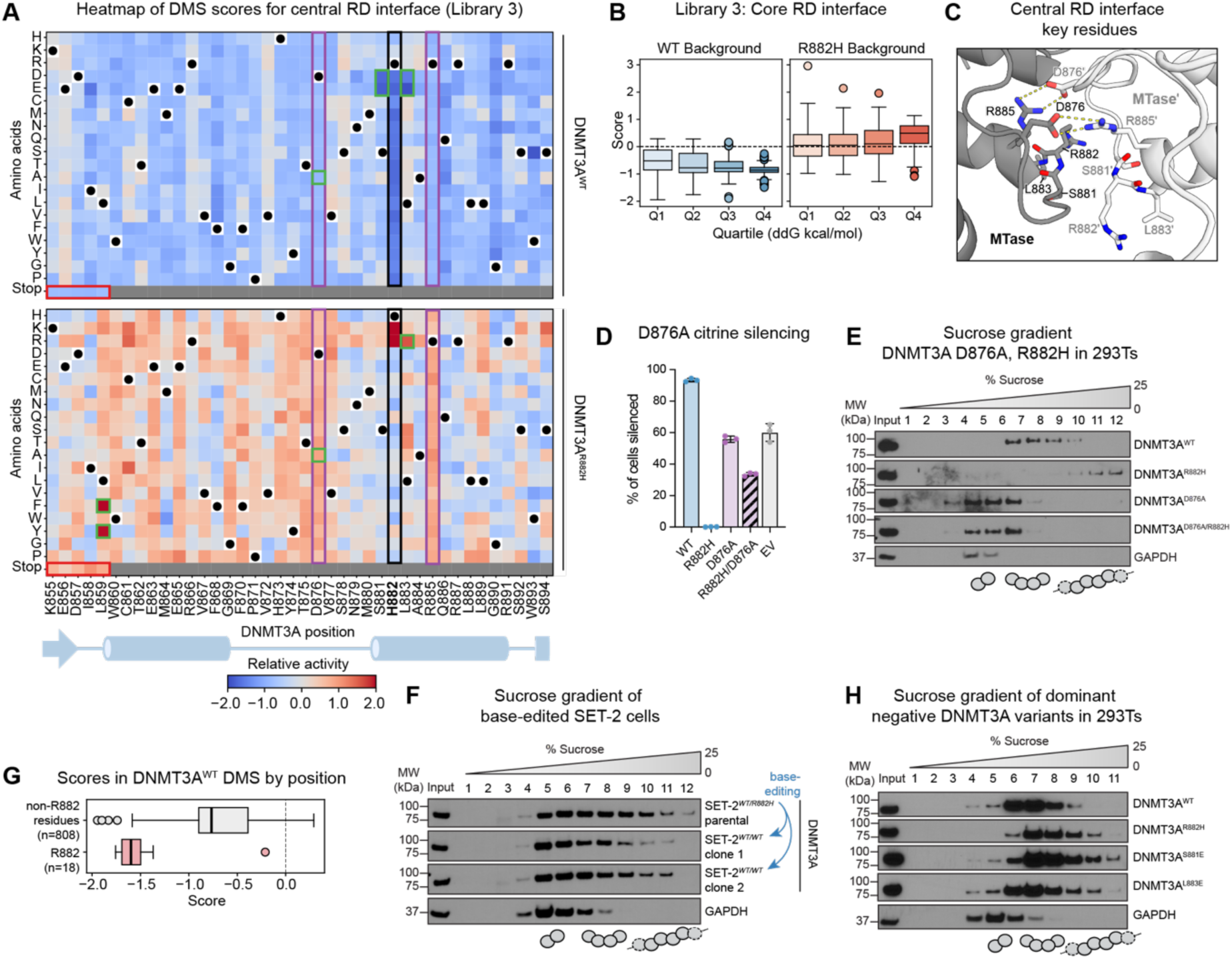
Disruption of the RD interface blocks aberrant macro-oligomerization by R882H. **A.** Heatmap showing activity scores of variants in deep mutational scanning of DNMT3A^WT^ (top) and DNMT3A^R882H^ (bottom) for the central RD interface library (Library 3). Color scale represents relative activity normalized to wild-type (top) or R882H (bottom). Black dots indicate wild-type residue. Black border indicates position 882, colored borders indicate residues and mutations of interest referenced in the text. **B.** Box plots of mutational scanning scores of each saturation library separated into quartiles based on ThermoMPNN predicted protein stability (Q1 most stable, Q4 least stable). center line, median; box, interquartile range; whiskers, up to 1.5 × interquartile range. See **Figure S6B** for Libraries 1 and 2. **C.** Closeup structural view of the RD-RD protein-protein interface showing the critical residues H873, D876, S881, R882, L883, and R885 in sticks. Monomer 1 in dark grey, monomer 2 in light grey. PDB: 8tdr. **D.** Flow cytometry quantification of reporter cells expressing indicated DNMT3A variants or empty vector (EV) with dox treatment after 21 days. Data are mean ± S.D. with individual replicates shown (n=3). **E.** Immunoblots of sucrose gradient ultracentrifugation experiment for indicated overexpressed FLAG-DNMT3A variants and GAPDH in HEK 293T cells. MW of DNMT3A2 monomer = 79kDa. MW of GAPDH tetramer = 148 kDa. **F.** Immunoblots of sucrose gradient ultracentrifugation experiment for DNMT3A and GAPDH from indicated SET-2 cell lines. **G.** Box plot showing scores of mutants at the R882 codon versus other positions in the 855-894 DNMT3A^WT^ library. Center line, median; box, interquartile range; whiskers, up to 1.5 × interquartile range. **H.** Immunoblots of sucrose gradient ultracentrifugation experiment for indicated overexpressed FLAG-DNMT3A variants and GAPDH. Data in **A, B** and **G** are mean of n = 3 biological replicates and the overall DMS experiment was performed once. Data in **E, F** and **H** are representative of n=2 independent experiments.

Notably, several missense mutants with negative enrichment scores in the wild-type-based library and positive enrichment scores in the R882H-based library were not previously shown or predicted to lead to loss of DNMT3A protein. In agreement, most of the clinical mutants located in the RD interface that were previously determined to be stable^35^ scored as less active than DNMT3A^WT^ in our deep mutational scan (**Figure S6C**). Notably, several of these mutants, including W860A, H873R/A, D876A, and N879A, have been previously shown to disrupt DNMT3A multimerization via the RD interface to generate dimers only mediated by the FF interface (**Figure 3C**)^10,11^. Of these, D876 and R885 involve the eponymous aspartate and arginine residues in the RD interface. While mutation of either of these residues in DNMT3A^WT^ caused loss-of-function, increased silencing in the DNMT3A^R882H^-based library was observed (**Figure 3A**, purple outline). As a representative of this category, we focused on D876A. Employing our fluorescent reporter, we confirmed that overexpression of DNMT3A^D876A^ provided no additional silencing activity over endogenous DNMT3A (**Figures 3D, S7A, and S7B**). By contrast, overexpression of DNMT3A^D876A/R882H^ still permitted reporter silencing by endogenous DNMT3A. Taken together, these results suggest that disrupting the protein-protein interaction between DNMT3A^R882H^ and endogenous DNMT3A suppresses the dominant-negative effect of R882H in a manner functionally equivalent to loss of DNMT3A^R882H^ protein altogether.

To investigate this behavior further, we sought to characterize the oligomeric state of DNMT3A and functional impact of these mutations by using purified proteins (**Figure S7C**). Profiling DNMT3A multimerization using size-exclusion chromatography and mass photometry (**Figures S7D–G**) revealed that the dominant species in DNMT3A^WT^ are tetramers, as anticipated^5,11^. By contrast, DNMT3A^D876A^ predominantly formed dimers, presumably with an intact FF and dissociated RD interface, while DNMT3A^R882H^ formed larger macro-oligomers. To assess DNMT3A multimerization in a cellular context, we used sucrose gradient ultracentrifugation to conduct size fractionation of DNMT3A overexpressed in HEK293T cells, using GAPDH as a size control (GAPDH tetramer MW = 148 kDa)^36^ (**Figure 3E**). This revealed that DNMT3A^R882H^ formed larger complexes than DNMT3A^WT^. By contrast, both DNMT3A^D876A^ and DNMT3A^D876A/R882H^ formed smaller complexes than DNMT3A^WT^, demonstrating that macro-oligomer formation promoted by R882H requires a binding-competent RD interface. To determine whether R882H can promote macro-oligomer formation at endogenous protein levels of DNMT3A, we conducted size fractionation in SET-2 cells, which are heterozygous *DNMT3A^WT/R^*^882^*^H^* ^37^ , as well as in SET-2 cells corrected to be *DNMT3A^WT/WT^* by base editing (**Figures 3F** and **S7H**). Consistent with our prior findings, we observed a higher proportion of higher molecular weight DNMT3A complexes in *DNMT3A^WT/R^*^882^*^H^* in comparison to *DNMT3A^WT/WT^*SET-2 isogenic cells, showing that R882H promotes macro-oligomer formation in an AML cancer cell line. Altogether, our results support the notion that R882H promotes macro-oligomerization and does not exert its dominant-negative effect by promoting dimer formation.

### R882 protects DNMT3A from macro-oligomerization

To investigate if macro-oligomerization drives the dominant-negative effects of R882H, we considered if other mutants may phenocopy this phenomenon. Notably, several mutants at the RD interface decreased DNMT3A^WT^ reporter silencing even more strongly than nonsense mutations (**Figures 1I and 3A**), consistent with dominant-negative behavior. These mutants are largely composed of R882 point mutations. Critically, mutation of R882 in DNMT3A^WT^ to almost any other amino acid scored as dominant negative, indicating that the dominant-negative effect of R882H is not due to the gain of histidine but rather the loss of arginine at position 882 (**Figures 3A and 3G**). This observation is consistent with mutations identified from patients with AML and clonal hematopoiesis, where not only R882H is observed but also R882C, R882P, and R882S^4,22^. In our deep mutational scan, the notable exception among R882 mutations was R882K, which scored nearly as active as DNMT3A^WT^. These data show the importance of maintaining a positively charged residue at position 882 for DNMT3A activity. Supporting this notion, L883R, which introduces positive charge just one residue away from position 882, partially rescued the activity in cis with DNMT3A^R882H^ (i.e., DNMT3A^R882H/L883R^) (**Figure 3A, green outline**); likewise, the octopus ortholog-inspired mutant DNMT3A^S878R/R882H/A884R^ also introduces positive charge and partially rescued activity (**Figure 1D**).

Notably, S881D/E and L883D/E also scored as dominant negative in the deep mutational scan; both mutations position a negatively charged amino acid directly adjacent to R882, potentially neutralizing the effect from its positive charge (**Figure 3B**). We considered if these adjacent mutations may phenocopy R882H and promote DNMT3A oligomerization. Sucrose gradient ultracentrifugation of DNMT3A^L883E^ and DNMT3A^S881E^ showed that both variants are biased towards forming larger species (**Figure 3H**), further suggesting that macro-oligomerization drives the dominant-negative phenotype. Together, our results demonstrate that maintenance of positive charge near the R882 position protects against dominant-negative inhibition of DNMT3A activity and protein macro-oligomerization.

### L859F rescues DNMT3A^R882H^ activity and oligomerization

Lastly, we considered the two rare, chemically similar mutations, L859F and L859Y, that could selectively rescue the activity of DNMT3A^R882H^ without significantly compromising that of DNMT3A^WT^. L859 is located near R882H, though they do not make direct contact (**Figure 4A**). Notably, introduction of either DNMT3A^L859F/R882H^ or DNMT3A^L859Y/R882H^ into our reporter system nearly fully restored enzyme activity, whereas DNMT3A^L859F^ and DNMT3A^L859Y^ did not greatly impact activity relative to DNMT3A^WT^, corroborating the scores obtained from the deep mutational scan (**Figures 4B, S8A, and S8B**). Strikingly, sucrose gradient ultracentrifugation demonstrated that DNMT3A^L859F/R882H^ formed smaller multimeric species than DNMT3A^R882H^, indicating that the rescue afforded by L859F may result from blocking aberrant macro-oligomerization (**Figure 4C**). In agreement, purified DNMT3A^L859F/R882H^ formed smaller species than DNMT3A^R882H^, as measured by size-exclusion chromatography and mass photometry (**Figures S7D and S8C–E**). Lastly, in a biochemical methyltransferase activity assay, DNMT3A^L859F/R882H^ displayed higher enzymatic activity in comparison to DNMT3A^R882H^ and DNMT3A^L859F^ (**Figure S8F**). In contrast to our deep mutational scan, DNMT3A^L859F^ was less active than DNMT3A^WT^ in the *in vitro* system, suggesting that the recruiting copy in the cellular reporting system, interaction with nucleosomes, or other factors in the cellular environment may influence the behavior of this mutant. Altogether, these data demonstrate that L859F rescues the dominant-negative effect of R882H and restores the tetrameric, active state of DNMT3A.

**Figure 4:**
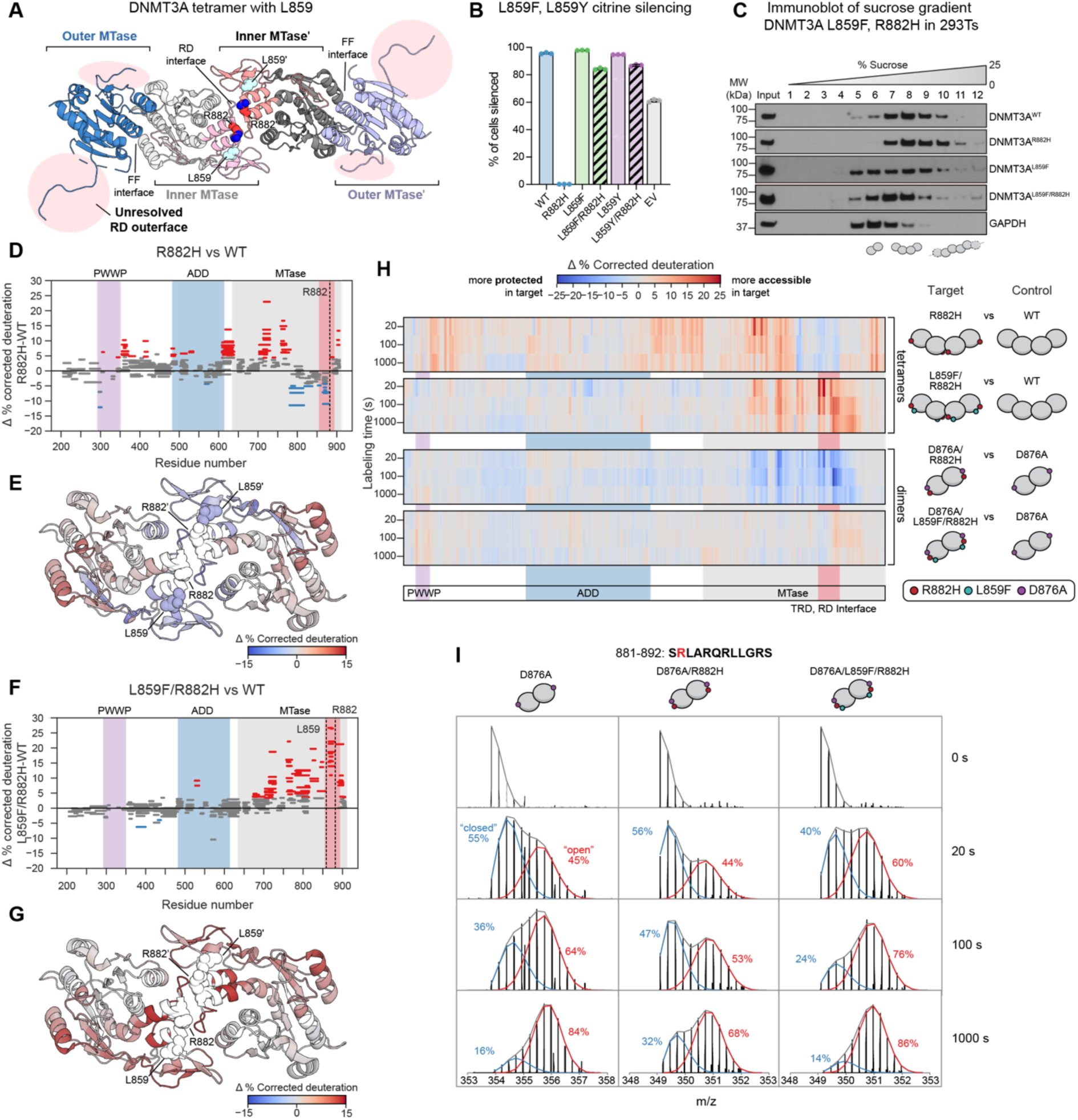
L859F rescues R882H activity and counteracts oligomerization. **A.** Structural view of the MTase tetramer with R882 (red and blue) and L859 (cyan) shown in spheres. The regions unresolved in the outer MTase protomers (blue) are highlighted with pink background and mapped to their corresponding regions on the inner MTase protomers (gray). PDB: 8tdr. **B.** Flow cytometry quantification of reporter cells expressing indicated DNMT3A variants or empty vector (EV) with dox treatment after 21 days. Data are mean ± S.D. with individual replicates shown (n=3). **C.** Immunoblot of sucrose gradient ultracentrifugation experiment for indicated overexpressed FLAG-DNMT3A variants and GAPDH. Data are representative of n=2 biological replicates. **D.** Woods plot showing deuterium differences (%) between WT and R882H tetramers at 100 sec. Deprotected (in DNMT3A^R882H^ relative to DNMT3A^WT^), protected, and non-significantly different peptides are shown in red, blue and grey respectively. Significant differences in % deuterium uptake at the peptide level were identified using a dual-threshold approach: (1) a two-tailed Welch’s t-test at α = 0.05, and (2) a difference threshold of 2 times the pooled standard deviation from % deuterium uptake measurements across all peptides. Additional Woods plots are shown in **Figures S9B, S9D, and S9E (Table S3)**. **E.** Differential deuterium uptake between R882H and WT at 100 sec mapped on the inner MTase (PDB: 8tdr). Per-residue coloring was determined using the Keppel and Weis algorithm^38^. **F.** Woods plot showing deuterium differences (%) between WT and L859F/R882H tetramers at 100 sec. Deprotected, protected and non-significantly different peptides are shown in red, blue and grey respectively. Significant differences in % deuterium uptake at the peptide level were identified using a dual-threshold approach: (1) a two-tailed Welch’s t-test at α = 0.05, and (2) a difference threshold of 2 times the pooled standard deviation from % deuterium uptake measurements across all peptides. **G.** Differential deuterium uptake between L859F/R882H and WT tetramer at 100 sec mapped on the inner MTase’ (PDB: 8tdr). Per-residue coloring was determined using the Keppel and Weis algorithm^38^. **H.** Heatmaps showing differential deuterium uptake within peptides across mutants at 20, 100, and 1000 sec. Only common peptides detected across all mutants are shown (**Table S4**). Mutant tetramers and D876A are normalized to wild-type tetramer; mutant dimers are normalized to D876A. Domain organization is indicated below. **I.** Representative isotopic distributions for peptide 881-892 across all time points for the dimeric variants. Binomial fitting to raw spectral data reveals bimodal isotopic distributions (99% confidence intervals). Binomial fits for the total population (gray), closed state (blue), and open state (red) are shown. Binomial fitting to raw spectral data for dimers and tetramers for peptides 865-880 and 881-892, are provided in **Figure S10**. Population fractions are shown as percentages. See also **(Table S5)**. Data are representative of n = 4 technical replicates. Data in **D**-**G** are average of n = 4 technical replicates

### R882H and L859F mutations alter pre-organization of the RD outerface

Having established that R882H promotes oligomerization and L859F restores the enzyme’s predominantly tetrameric state, we next sought to determine how this rescue occurs and whether it could shed light on the mechanism of R882H itself. As recent X-ray crystal structures of the wild-type and R882H tetramers are nearly identical (**Figure 1B**, <1 Å R.M.S.D.), we hypothesized that R882H and L859F might instead change the conformational dynamics of the RD interface region. This hypothesis was in part based on the fact that, in contrast to the stable RD-RD interface, the solvent-exposed RD residues of the outer protomers (which we refer to as the “outerface”) could not be resolved (**Figure 4A**, amino acids 711-725, 798-832, 846-890, in pink)^8^, supporting the notion that this region may be dynamic and flexible when not bound to another copy of DNMT3A. Notably, L859 is part of the differentially folded RD interface region and, like R882H, was not resolved in the outerface.

To interrogate the dynamics of the RD interface residues and the broader enzyme, we performed HDX-MS on SEC purified full-length tetramers of DNMT3A^WT^, DNMT3A^R882H^, and DNMT3A^L859F/R882H^ protein variants (**Figures 4D-I and S9A**). Experiments were conducted at low temperature to slow exchange kinetics and capture dynamics in highly flexible regions over early time points using our recently developed nanoflow configuration (nHDX), which enabled comprehensive analysis with minimal protein consumption. Decreased levels of deuterium uptake were observed for peptides at the RD interface in DNMT3A^R882H^ in comparison to DNMT3A^WT^ across all timepoints (**Figures 4D, 4E, and 4H**) suggesting that the RD interface region of DNMT3A^R882H^ is less accessible than that of DNMT3A^WT^. In addition, we observed increased accessibility in both the PWWP domain and the ADD-MTase linker in DNMT3A^R882H^ versus DNMT3A^WT^ (**Figure 4D**), suggesting broader movement of the regulatory domains. Strikingly, introduction of L859F broadly increased deuterium uptake across the MTase domain in DNMT3A^L859F/R882H^ to levels even higher than that observed for DNMT3A^WT^, rescuing the decreased accessibility observed for DNMT3A^R882H^ (**Figures 4F and 4G)**. L859F also suppressed the increased accessibility observed in the PWWP domain and the ADD-MTase linker of DNMT3A^R882H^, as DNMT3A^L859F/R882H^ exhibited similar levels of deuterium uptake as DNMT3A^WT^. Altogether, these data suggest that the R882H mutation causes striking changes in protein dynamics, notably rigidifying the RD interface region, and that these effects can be reversed by L859F. Our HDX-MS data further reveal that the RD region exists in a conformational equilibrium between a dynamic, fast-exchanging ‘open’ state and a stable, slow-exchanging ‘closed’ state. The R882H mutation shifts this population toward the closed conformation, an effect that is reversed by the L859F/R882H mutation, which restores the open state (**Figure S10**).

Due to our explicit use of tetramers, to directly test whether changes in deuterium uptake among the DNMT3A variants were driven by the inner or outer protomers or both, we introduced the D876A mutation to disable the bound RD interface and conducted nHDX-MS on the purified dimers to explicitly analyze the unbound state. As expected, the DNMT3A^D876A^ dimer showed increased accessibility across RD residues compared to the wild-type tetramer (**Figures S9B and S9C**). Consistent with our hypothesis, the R882H mutation in DNMT3A^D876A/R882H^ induced large-scale protection across the MTase domain relative to DNMT3A^D876A^ (**Figure 4H**). Strikingly, this protection was completely abrogated by the L859F mutation, as DNMT3A^L859F/D876A/R882H^ displayed deuterium uptake nearly identical to the DNMT3A^D876A^ dimer, demonstrating that L859F rescues the rigidification caused by R882H. Analysis of the exchange kinetics revealed that while the DNMT3A^D876A^ and DNMT3A^D876A/R882H^ show a dynamic interconversion between open and closed states, DNMT3A ^L859F/D876A/R882H^ exists predominantly in the open state from the earliest time points measured (**Figure 4I and S10**).

Taken together, these findings are consistent with a dynamic RD region, as suggested by the unresolved outerface in the crystal structure and our experiments on both oligomeric states, and show that the R882H mutation stabilizes a protected, rigid conformation in the RD region. Interestingly, changes in the deuterium uptake in the N-terminal regulatory domains and disordered regions are not observed in the DNMT3A dimer, showing that formation of the bound RD-RD interface in tetramers is coupled to significant conformational dynamics of the broader protein, including the regulatory domains **(Figure 4H)**. Altogether, these data therefore support a model in which pre-organization of the dynamic RD outerface into a binding-competent state promotes DNMT3A oligomerization, and this process is enhanced by the hotspot R882H mutation, ultimately inhibiting the enzyme’s activity. The corresponding restoration of conformational dynamics at the unbound RD outerface by L859F provides the mechanistic basis for the rescue of the enzyme’s activity and proper tetramer formation (**Figure 5**).

**Figure 5:**
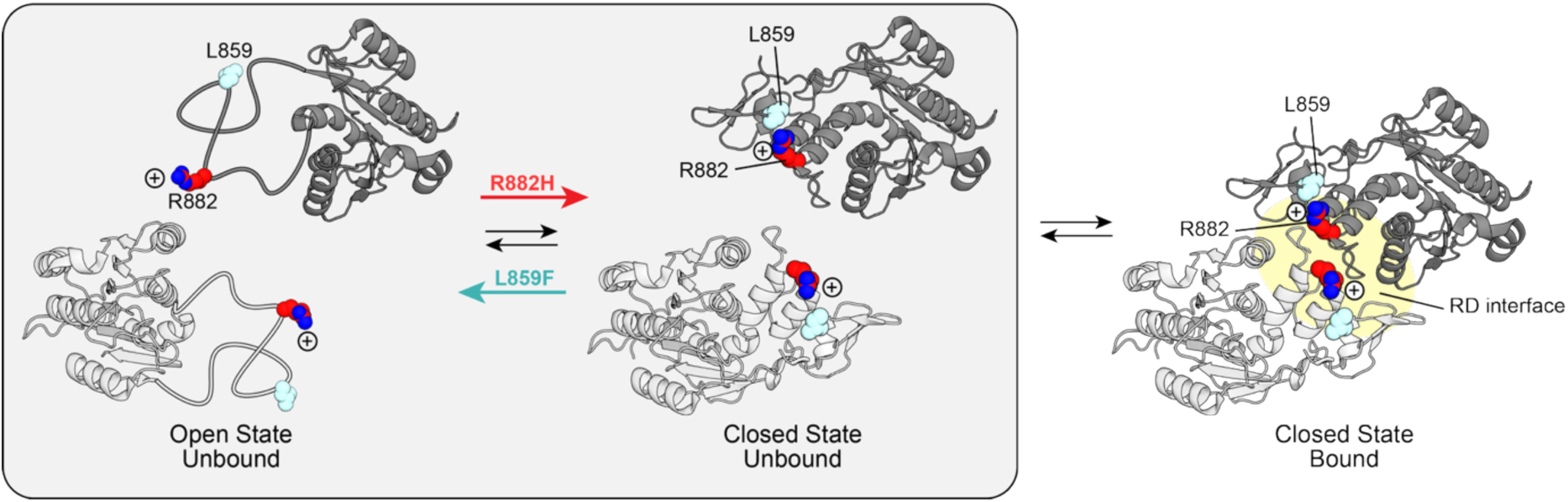
Cartoon model of R882H and L859F influence on the RD outerface equilibrium between open and closed states and subsequent interface formation. The RD interface residues of DNMT3A are in equilibrium between an open and closed state; R882H promotes the closed state, and L859F promotes the open state. The population of closed, unbound state can form RD-RD interfaces. R882H is colored in red (carbons) and blue (nitrogen), L859F in cyan, and RD-RD interface highlighted in yellow. PDB: 8tdr.

## Discussion

The *DNMT3A R882H* hotspot mutation is one of the most prevalent somatic mutations in humans, yet the biochemical mechanism underlying its dominant-negative activity has remained controversial. Here, we implemented paired deep mutational scanning to chart the genetic landscape surrounding the DNMT3A R882H hotspot. Deeper analysis and investigation of these diverse mutations reveals numerous mechanisms that control DNMT3A oligomerization and activity through the TRD and the core RD interface. We show that DNMT3B paralog-informed mutations in the TRD tune the DNA binding and autoinhibition of both DNMT3A^WT^ and DNMT3A^R882H^, consistent with the different sequence preferences and genomic roles of DNMT3A and DNMT3B. In the core RD interface region, we discover a dominant-negative hotspot that reveals the critical role of positive charge around position R882 in protecting DNMT3A from aberrant macro-oligomerization. Although we do not rule out that the gain of histidine from R882H directly enhances RD-RD protein-protein interactions, our results strongly support a model in which loss of arginine at the critical R882 safeguarding position primarily drives macro-oligomerization, consistent with clinical data showing that other diverse R882 mutations also lead to clonal hematopoiesis^22^. This loss-of-function mechanism was likely overlooked, since it is not obvious from prior crystal structures, highlighting the value of using unbiased, massively parallel mutational scanning for uncovering and parsing protein functional hotspots.

Finally, we study wild-type, R882H and unexpected rescue mutations with HDX-MS, a powerful approach providing insights into protein dynamics that are inherently challenging to capture with static structural methods like X-ray crystallography. We discover that R882H pre-organizes the unstructured RD outerface, supporting a model in which rigidification of the RD outerface is a key biochemical driver of aberrant DNMT3A macro-oligomerization. Restoration of conformational flexibility in this region by L859F rescues the enzyme’s activity by preventing oligomerization. Taken together, our results suggest that positive charge at position 882 prevents pre-ordering, although the exact mechanism warrants further study.

More broadly, we reveal that protein conformational dynamics is an essential factor underlying the mechanism of DNMT3A^R882H^ and likely endogenous regulation of the enzyme, as we observe that the formation of the RD-RD interface is linked to larger movements in the regulatory domains. We anticipate these unexpected findings will open new avenues for future investigation. Collectively, our study further highlights how protein dynamics play critical roles in mediating protein-protein interactions and molecular recognition and examines the striking mechanism of a disease-associated mutation enhancing a protein-protein interface in high molecular detail. Protein-protein interactions are frequently perturbed by mutations in human disease^39,40^. Point mutations can alter the interactions between protein subunits in homo-multimers, which are highly sensitive to genetic variation and lie at the verge of supramolecular self-assembly^24,41^ Our observation that R882H results in loss of a safeguard, rather than directly creating an enhanced interface^42^, matches prior work showing that negative design elements in protein-protein interfaces are strongly evolutionarily conserved to safeguard against self-assembly^1,24,43^. Although single point mutations that disrupt these safeguards have been widely documented to drive self-assembly in many proteins^24^, our work provides the first example of this mechanism involving a cancer mutation to aberrantly promote protein-protein interactions. Notably, homo-multimers are predicted to involve 20% of the human proteome^41^, raising the prospect that other human cancer mutations might reprogram protein-protein interactions through a similar mechanistic paradigm as the R882H hotspot mutation. In summary, we unveil the mechanism of the R882H hotspot mutation, deepening our understanding of protein-protein interfaces and how they are subverted in human disease.

## Materials and Methods

### Cell culture

The following cell lines were used: K562 (ATCC), HEK293T (a gift from B.E. Bernstein, Massachusetts General Hospital), SET-2 (a gift from M.D. Shair, Harvard University), Massachusetts General Hospital). All cell lines were cultured in a humidified 5% CO_2_ incubator at 37℃ and were tested for mycoplasma. K562s were cultured in RPMI-1640 (Gibco) with 10% FBS and 100 U ml^−1^ penicillin and 100 μg ml^−1^ streptomycin (Gibco). HEK293Ts were cultured in DMEM (Gibco) with 10% FBS and 100 U ml^−1^ penicillin and 100 μg ml^−1^ streptomycin (Gibco). SET-2s were cultured in RPMI-1640 (Gibco) with 20% FBS and 100 U ml^−1^ penicillin and 100 μg ml^−1^ streptomycin (Gibco). FBS was obtained from Atlas Biologicals.

### Lentiviral Production

To produce lentivirus, transfer plasmids were co-transfected with GAG/POL and VSVG plasmids into HEK293T cells using Lipofectamine 3000 (Thermo Fisher Scientific) according to the manufacturer’s protocol. The medium was exchanged 6-8 hours after transfection. After 48-60 hours, the medium was collected, passed through a 0.45 μm filter, and either used immediately or snap frozen and stored at -80℃ until use.

### Lentiviral Transduction

K562 and SET-2 cells were transduced by spinfection (1,800xg, 90 min, 37℃) with 12 or 8 μg/mL polybrene (Santa Cruz Biotechnology), respectively. HEK293T cells were transduced by incubation with the filtered viral medium and 10 μg ml^−1^ polybrene for 48-72 hours, after which the medium was replaced. Transduced K562 and Set2 cells were selected using 2 μg ml^−1^ puromycin (Gibco) or FACS. Transduced 293T cells were selected with 1 μg ml^−1^ puromycin (Gibco).

### General Plasmid Construction

All plasmids except base-editing plasmids were cloned by Gibson Assembly using NEBuilder HiFi (New England Biolabs). Cloning strains used were NEB Stable (lentiviral vectors) and NEB 5-alpha (other plasmids) (New England Biolabs). Bacterial cultures were grown at either 30℃ (for lentiviral vectors) or 37℃ (other plasmids). For base editing plasmids, sgRNAs were ordered as synthetic oligonucleotides (Sigma-Aldrich), annealed and ligated into pRDA_479, (Addgene #179099). Final constructs were validated by Nanopore and/or Sanger sequencing (Plasmidsaurus and Quintara Biosciences). Plasmids and oligonucleotides are provided in **Table S6.**

### Reporter cell line generation

The reporter cell line used for individual variant timecourse silencing assays was previously reported in ref^14^. This cell line was further modified into the DNMT3A knock-out cell line using a CRISPR RNP. All CRISPR reagents were obtained from Integrated DNA Technologies (IDT) unless otherwise specified. To knock out *DNMT3A* in our reporter cell line, we used our previously reported DNMT3A KO sgRNA sgNL5 (see guide sequence in **Table S6**). The guide RNA was ordered as an Alt-R CRISPR-Cas9 crRNA. To prepare the gRNA complex, crRNA and Alt-R CRISPR-Cas9 tracrRNA were mixed in equimolar ratio, heated to 95 °C for 5 min, and then allowed to cool to room temperature. To prepare the RNP complex, the sgRNA complex was combined with the Alt-R Cas9 enzyme and incubated at RT for 20 min. The Neon Transfection

System was used to electroporate 2 × 10^5^ *DNMT3A*^WT^ reporter cells with the assembled RNP complex. Electroporated cells were recovered in antibiotic-free media for 72 h. Single cell clones were sorted by FACS and screened by western blot.

For the DMS reporter cell line, the PuroR-9xTet0-pEF-IGKleader-hIgG1_FC-Myc-PDGFRb-T2A-Citrine-PolyA reporter cassette from Addgene Plasmid 161927 ^28^ was cloned into a lentiviral vector pSPS2 and transduced into K562 cells. These cells were then transduced with pEF-H2B-mCherry-T2A-rTetR-dDNMT3A2 Several mCherry and citrine positive clones were sorted and one clone was selected for use in the DMS.

### Reporter silencing assays

Reporter silencing assays were conducted as described in ref^26^ with the following changes: cells were treated with 1 μg ml^−1^ dox (Sigma-Aldrich), and gates were set based on reference reporter cells cultured in parallel without dox and reference fully silenced reporter cells (See gating scheme in **Figure S3D**). Briefly, cells were transduced with the appropriate lentiviral vector expressing a DNMT3A variant, selected with 2 μg ml^−1^ puromycin (Gibco) for 5-7 days, subjected to 2-3 days of puromycin washout, then plated for dox treatment. Cells were subsequently passaged and measured for silencing every 2-3 days, maintaining cell count <1.2 M cells ml^−1^.

### Protein expression and purification

Full-length DNMT3A2 was recombinantly expressed and purified as described in ref^26^ with the following modifications: LB medium included 50 μg ml^−1^ kanamycin and 25 μg ml^−1^ chloramphenicol, two washes with 50 mL of base buffer containing 100 mM imidazole were performed before elution with base buffer containing 200 mM imidazole. The eluate was concentrated using an Amicon® Ultra Centrifugal Filter, 30 kDa MWCO and exchanged into DNMT3A storage buffer consisting of 5 mM HEPES pH 8.0 cold, 75 mM NaCl, and 0.25 mM TCEP with an Econo-Pac 10DG Desalting Column (BioRad). The protein was injected onto a Superose 6 Increase 10/300 GL column (Cytiva), eluting in storage buffer. Eluate was concentrated using the same method as described for concentration.

### 3H Activity assays

DNMT3A activity was measured as described in ref^14^ with the following modifications: storage buffer was diluted before beginning experiments to final concentrations of 20 mM HEPES pH 8.0 cold, 300 mM NaCl, and 1 mM TCEP. 0.2μM DNMT3A2 was used, and reactions were incubated at room temperature for 3 hours. For histone stimulation experiments, the amount of H3K4me0 peptide (ARTKQTARKSTG-NH2, Biomatik) added is specified in each figure legend.

### Sucrose gradients

Cell pellets containing 2 million cells were thawed on ice, then resuspended at 50 μL/1 million cells in RIPA buffer supplemented with 1x Halt Protease Inhibitor Cocktail, 1 mM PMSF, and 2 mM EDTA and incubated on ice for 30 minutes. Then, the lysate was supplemented with an equivalent volume of wash buffer (50 mM Tris-HCl pH 8, 2mM MgCl_2_, 50 mM NaCl) containing 1x Halt Protease Inhibitor Cocktail, 1 mM PMSF, and 500 U/mL Benzonase Nuclease (Millipore Sigma) and incubated while rotating at room temperature for 20 minutes. The lysate was centrifuged at 18,000xg for 10 minutes at 4℃. The supernatant was transferred to a fresh tube and labeled as input. 5% and 25% w/v sucrose solutions were prepared with final concentrations of 50 mM Tris pH 8.0 cold, 300 mM NaCl, and 1 mM TCEP. A BioComp Gradient Fractionator was used to make 5-25% sucrose gradients in 7/16 x 2-3/8 Centrifuge Tubes (Beckman Coulter). 150 μL of input was loaded onto the top of the sucrose gradient and the tubes were loaded into a SW60Ti rotor. The samples were centrifuged in a Beckman Coulter Optima L-90K at 50,000 rpm for 6 hours at 4℃. 300 μL fractions were collected using the BioComp Gradient Fractionator, flash frozen, and stored at -80℃ until immunoblotting was performed.

### Immunoblotting

Samples were loaded onto a Bolt 4-12% Bis-Tris Plus WedgeWell gel (Invitrogen), electrophoresed, and transferred onto a PVDF membrane using the GenScript eBlot. Membranes were blocked in Tris-buffered saline Tween (TBST) with 5% BSA for one hour. Membranes were incubated at 4℃ overnight with primary antibody diluted in TBST with 5% BSA: DNMT3A (Cell Signaling Technology, catalog no. 32578, D2H4B, 1:5000), FLAG (Sigma-Aldrich, catalog no. F1804, M2, 1:2,000), GAPDH (Santa Cruz Biotechnology, catalog no. sc-47724, 0411, 1:8,000). Membranes were washed 3x with TBST and incubated with secondary antibody for 1 hour at room temperature: anti-mouse IgG HRP conjugate (Promega, catalog no. W4021, 1:40,000 for FLAG, 1:400,000 for GAPDH), anti-rabbit IgG HRP conjugate (Promega, catalog no. W4011, 1:100,000 for DNMT3A). Membranes were washed 3x with TBST and imaged using SuperSignal West Pico PLUS or SuperSignal West Femto chemiluminescent substrates (Thermo Fisher Scientific).

For immunoblotting of SET-2 sucrose gradients, 300 μL fractions were precipitated with 100% (w/v) Tri-chloro acetic acid (TCA), washed 2x with 100% acetone, and reconstituted in 50 μL 80% PBS+ 18% NuPAGE™ LDS Sample Buffer (Invitrogen) + 2% 2-Mercaptoethanol (Sigma) before loading onto a Bolt 4-12% Bis-Tris Plus WedgeWell gel (Invitrogen) and proceeding as with all other immunoblots.

### Library generation for deep mutational scanning

For each sub-library (aa 660-685, 815-854, 855-895) a separate destination vector was generated using Gibson Assembly (DEST1a, DEST2a, and DEST3 in **Table S6**) based on pEG1.175A with a BamHI restriction site in place of the library fragment. For libraries 1 and 2, secondary destination vectors DEST1b and DEST2b were also generated with the R882H mutation. (See **Figure S3C**) A custom python script (see **Supplemental Code**) was used to generate the saturation and DNMT3B informed library sequences. For the saturation library, codons were chosen to be 2 or 3 nucleotides WT DNA sequence if possible, and then prioritized by human codon frequency. For the DNMT3B library, the DNMT3B codon was used. After generating these sequences, appropriate overhangs and barcodes were added, to enable amplification of sub libraries from the full set of sequences, and the library was ordered as a Twist oligo pool. Each sub library was amplified from the total pool with 15 cycles of PCR (NEBNext Ultra II Q5 Master Mix) targeted to the barcodes, followed by 10 cycles of amplification with primers targeting the Gibson Assembly overhangs to remove the barcodes. Finally, sub libraries were assembled into the appropriate destination vector using NEBuilder HiFi (New England Biolabs). Completed reactions were isopropanol-precipitated, resuspended in 6 μL of H_2_O and 1.5 μL was electroporated into Lucigen Endura Electrocompetent cells. After overnight growth on LB-agar plates with 100 μg/mL ampicillin, coverage of each library was confirmed to be >1000 cfu per library variants by plating of serial dilutions. Plates were scraped and DNA was isolated via MaxiPrep (ZymoPURE II Plasmid MaxiPrep Kit) to generate the final plasmid libraries.

### Deep mutational scanning

Lentivirus was generated as described above for each of the six libraries independently. Prior to large-scale infection for screening, lentiviruses were titered by measuring transduced cell counts after puromycin selection. Reporter cells were infected with lentivirus with a multiplicity of infection of <0.4 and at least 700x coverage of the library, then selected with puromycin for 5 days. Following selection, each pool of cells was split into three replicates and treated with 0.1 μg/mL of dox for 4 days (WT background libraries) or 14 days (R882H background libraries) to target sorting to a timepoint with 10-40% of cells silenced. Cells were sorted on a FACSAria II (BD) or FACSAria Fusion, collecting citrine+, citrine–, and unsorted (all mCherry+) cells. Fully silenced cells were used as a control for citrine gating (see **Figure S3D** for full gating scheme).

### Library preparation and sequencing

To prepare for sequencing, genomic DNA was isolated using the QIAamp DNA Blood Mini kit. All samples maintained at least 100x library coverage after gDNA extraction. 20 cycles of PCR with targeted primers (**Table S6**) and NEBNext Ultra II Q5 Master Mix with 1000 ng DNA per 50 uL reaction was used to amplify the region of interest for each library. This was followed by a second round of amplification (8 cycles) with barcoded primers, and finally sequencing on an Illumina NextSeq1000.

### Data analysis

Raw paired-end sequencing reads were trimmed with cutadapt (v4.9)^44^ and merged with bbmerge (v39.06)^45^. Then, Enrich2 was used to calculate variant counts and estimate log enrichment scores and errors^29^. The WT library was normalized to DNMT3A^WT^, and the R882H library was normalized to DNMT3A^R882H^. Only sequences with greater than 50 reads in the day 0 and unsorted conditions were used in downstream analyses. Information from Enrich2 output was further merged into **Table S1**. Library 2 WT replicate 2 was excluded from further analysis due to poor correlation with replicates 1 and 3, but replicate correlation plots reflecting this replicate are still included in **Figure S4** and the processed data is included in **Table S1**. Jupyter notebooks for plotting heatmaps, scatterplots, and other visualizations can be found in **Supplemental Code**.

### Combinatorial mutation data analysis

Custom python code (see **Supplemental Code**) was used to perform combinatorial mutational analysis. Statistics were calculated using scipy (v1.8.1). Linear regression was performed using sklearn (v0.0.post9). Linear regression parameters in **Figure 2C** correspond to the equation:

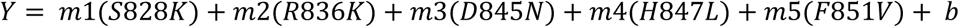

Where Y=variant score, m1:m5 are the coefficients to be fit, and b is the y-intercept.

### Fluorescence polarization

Purified DNMT3A2 protein was diluted to 5 μM in storage buffer (20 mM HEPES pH 8.0, 300 mM NaCl, 1 mM TCEP) then diluted 1:5 into assay buffer (50mM Tris-HCl pH 8.0, 1mM TCEP, 0.1mg/mL BSA), containing 10 nM Cy3-labeled 30-bp DNA probe^11^ (**Table S6**) and 1 or 10 μM H3K4me0 peptide (ARTKQTARKSTG-NH2, Biomatik). The protein-probe mixture was distributed into wells of a black 384-well plate (Corning) in triplicate. Then, a twofold serial dilution was performed in assay buffer with 10 nM labeled DNA probe and 1 or 10 μM H3K4me0 peptide for a final volume of 20 μL per well. The plate was incubated in the dark at room temperature for 90 minutes and read (1,700-ms integration) using a SpectraMax i3x with a rhodamine fluorescence polarization cartridge and SoftMax Pro software (Molecular Devices). The G-factor (3.1) was adjusted to set the polarization of assay buffer and 20 nM probe only to a reference value of 37 mP. Curves were fit with a constant DNA concentration of c=10 nM using scipy( v1.8.1) to the equation:

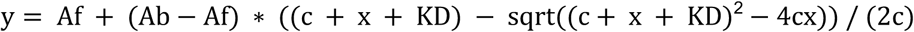

where (y) the measured signal is dependent on the contributions of bound (Ab) and free (Af) fluorescently labeled DNA as a function of total protein concentration (x) and ligand concentration (c = 10nM). FP signal measurements at 1 μM protein concentration were compared with unpaired, two-tailed T tests performed in GraphPad Prism (v.10.4.0).

### Mass photometry

A Refeyn TwoMP mass photometer equipped with AcquireMP(version 2024 R1.1) was used to collect mass photometry data. Briefly, purified DNMT3A2 proteins were diluted to 2 μM in storage buffer (20 mM HEPES, 1mM TCEP, 300 mM NaCl), left at 4C overnight, then diluted to 200 nM protein (3 times, once for each replicate) in storage buffer ∼1 hour before measurement. The instrument was allowed to warm up for 1 hour prior to calibration and measurement. No. 1.5H high precision glass coverslips (24×50 mm), (Thorlabs CG15KH) were cleaned 3x with ultrapure water and isopropanol and dried under nitrogen gas, then affixed with a reusable silicone gasket. Zeiss Immersol 518 F immersion oil was added to the objective, and the slide was secured to the instrument. To calibrate, 18 μL of assay buffer was placed on the slide and the instrument was focused. Then, 2 μL of 10x Calibration mix (0.03 µM Thyroglobulin 0.1 µM BSA in PBS and 0.05% glycerol) was added to the droplet and mixed with a pipette set to 18 uL. Video acquisition was performed for 1 minute. The same procedure was used to measure each DNMT3A2 variant in triplicate, with a final concentration of 20 nM. DiscoverMP (version 2024 R1) was used to conduct preliminary data analysis (generating a calibration curve and calculating masses from measured events), and custom python scripts (**Supplemental Code**) were used to assign masses to multimeric states of DNMT3A2 using a gaussian mixture model (sklearn v0.0.post9).

### SET-2 base editing

Two manually designed sgRNAs (**Table S6**) were cloned into the lentiviral base editor (ABE8e) vector pRDA479 (Addgene 179099) by Golden Gate Assembly. Lentivirus was generated and SET-2 cells were spinfected and selected as described above. After selecting one of the two sgRNA edited pools of cells based on a higher editing rate (sgA3), single clones were sorted, screened with Sanger sequencing, and confirmed by Illumina sequencing for cells that were WT/WT at the R882 position (as opposed to WT/R882H found in unedited SET-2).

### Phylogenetic analysis

To explore the phylogeny of DNMT3A, an initial set of homologous sequences was collected using HMMER ^46^ (v3.4) searched with the hsDNMT3A sequence (uniprot id Q9Y6K1). From this initial set of sequences, 10 sequences containing all three domains in DNMT3A (PWWP, ADD, MTase) were selected from which a profile HMM was generated, prioritizing well validated model organisms as well as diversity of animal clade representation (human - Q9Y6K1, mouse - O88508, chicken - Q4W5Z4, zebrafish - Q588C2, octopus - A0A0L8HNR6, parasitic nematode - A0A0V0S0Y6, japanese rice fish - A0A3B3H694, alligator - A0A3Q0HD35, honeybee - A0A7M7MWZ4, starfish - A0A8B7YM53). The profile HMM was built by aligning the above sequences using MAFFT with the E-INS-i setting then running HMMbuild with the default parameters. This profile was then used with HMMsearch to search the UniProtKB database with a relative bitscore threshold of 0.5. HMMalign was then used to align the resulting sequences.

After making the preliminary alignment, the sequences were filtered for those that were at least 600 amino acids in length, to identify sequences long enough to contain all three DNMT3 domains. Then, the filtered alignment was realigned with Clustal Omega (v1.2.4)^47^. Then, this alignment was used to make a phylogenetic tree with FastTree (v2.1.11)^48^ which was further analyzed and visualized using custom python scripts and packages bioviper (v0.1.2)^49^ , logomaker (v0.8)^50^, ete3 (v3.1.3)^51,52^ , biopython (v1.78), code adapted from ^51^ and iToL (v7)^53^.

### HDX-MS sample preparation

The HDX-MS analysis was performed on a recently described nanoflow HDX-MS configuration (nHDX-MS) consisting of an automated sample handling robot (CHRONECT™ HDX PAL Parallel Extended, Trajan Scientific and Medical), Ultimate™ 3000 UHPLC, and Orbitrap Exploris™ 480 Biopharma Edition (Thermo). Briefly, 3 μL DNMT3A constructs (50 ng/μL in 20mM HEPES, 300mM NaCl, 5mM TCEP) were diluted with 15 μL D2O buffer (20mM HEPES, 300mM NaCl, 5mM TCEP) at pH 7.5 to initiate the deuterium exchange. Samples were quenched at 20, 100, and 1000 sec with addition of 45 μL of cold guanidine hydrochloride (4 M, 0.5% formic acid, 5 mM TCEP). Samples were digested online for 3 min at 200 μl/min using an immobilized protease type XIII/pepsin column (w/w, 1:1, NBA2014002, NovaBioAssays), operated at 8 °C. Peptides were trapped onto an Acclaim PepMap™ 100 µ-Precolumn (C18, 0.3mm × 5mm, 5µm, Thermo) using solvent A (0.23% formic acid) and were separated onto a PepSep C18 (0.15mm × 150mm, 1.9 µm, Bruker) at 1 μl/min with a gradient of 10% to 30% solvent B (0.23% formic acid in acetonitrile) applied over 12 minutes. Both the trap and analytical columns were housed in a temperature-controlled box at 0 °C. Peptides were introduced into mass spectrometry using an EASY-Spray™ Source coupled with EASY-Spray™ emitter (15 µm ID, Thermo). The spray voltage and transfer capillary temperature were set to 1.9 kV, and 250 °C, respectively. For peptide identification, samples were analyzed in data-dependent acquisition mode. Full MS scans were acquired in the 300 to 900 Th range, with a resolution of 120K at 200 m/z, and RF lens set at 40%. MS/MS acquisition parameters were as follows; resolution of 7.5K at 200 m/z, normalized AGC target of 30%, HCD collision energy of 30, max fill time of 50 ms, RF lens at 40%, and isolation window of 0.7 m/z. Unassigned and singly charged ions were excluded from MS/MS. Data were acquired in profile mode in MS and centroid mode in MS/MS scans. For HDX-MS measurements, samples were analyzed in data-independent acquisition (DIA) mode with an interscan method (https://pubs.acs.org/doi/full/10.1021/acs.analchem.5c00429).ach acquisition cycle started with a full MS scan in the 300 to 920 Th range, with resolution of 60 K at 200 m/z, and RF lens set at 40%. This was followed by a DIA MS/MS scan from 300 to 900 Th, and a subsequent DIA MS/MS scan from 310 to 910 Th with isolation width of 20 Th. The window overlaps in each DIA MS/MS scan were set to 0 Th. The DIA MS/MS acquisition parameters were as follows; resolution of 30K at 200 m/z, normalized AGC target of 500%, HCD collision energy of 27, max fill time of 54 ms, and RF lens at 40%.

### Peptide Identification using Spectrum Mill

The peptide identification runs were searched using SpectrumMill Proteomics Workbench (prerelease version BI.08.02.218, Agilent Technologies). A non-specific enzyme search was performed using a SwissProt HUMAN FASTA file (release date January 16, 2020) including DNMT3A mutants, a set of common contaminant proteins (264 sequences), pepsin and protease type XIII. Peptide and fragment tolerances were at ±20 ppm, protein and peptide false discovery rate at 1%, and minimum matched peak intensity at 40%.

## HDX-MS data analysis

The HDX-MS runs processed using HD-Examiner™ Pro application in CHRONECT Studio (ver. 0.9.0.104, Trajan Scientific and Medical). DNMT3A peptide features (Sequence, m/z, and retention time) identified from the peptide identification runs were used as input for processing the DIA data. Spectra were allowed ±10 ppm mass tolerance for precursor and product ions. The retention time (RT) integration window was set to 0.2 min with RT variance of 0.4 min. The extracted ion chromatograms (XICs) were smoothed using Savitsky Golay smoothing (window size of 5). The following peptide quality filters were used to compare the deuterium uptake across different DNMT3A mutants across timepoints; minimum number of fragments set to 3, and maximum % deuterium standard deviation of 5. For comparison of deuterium uptake across all mutants, minimum number of fragments was set to 2, and maximum % deuteration standard deviation was set to 7. The deuterium content of the peptides was calculated assuming loss of 2 Da from the N-terminus of the peptides (N-P-2 enabled).

---

**Figure S1:**
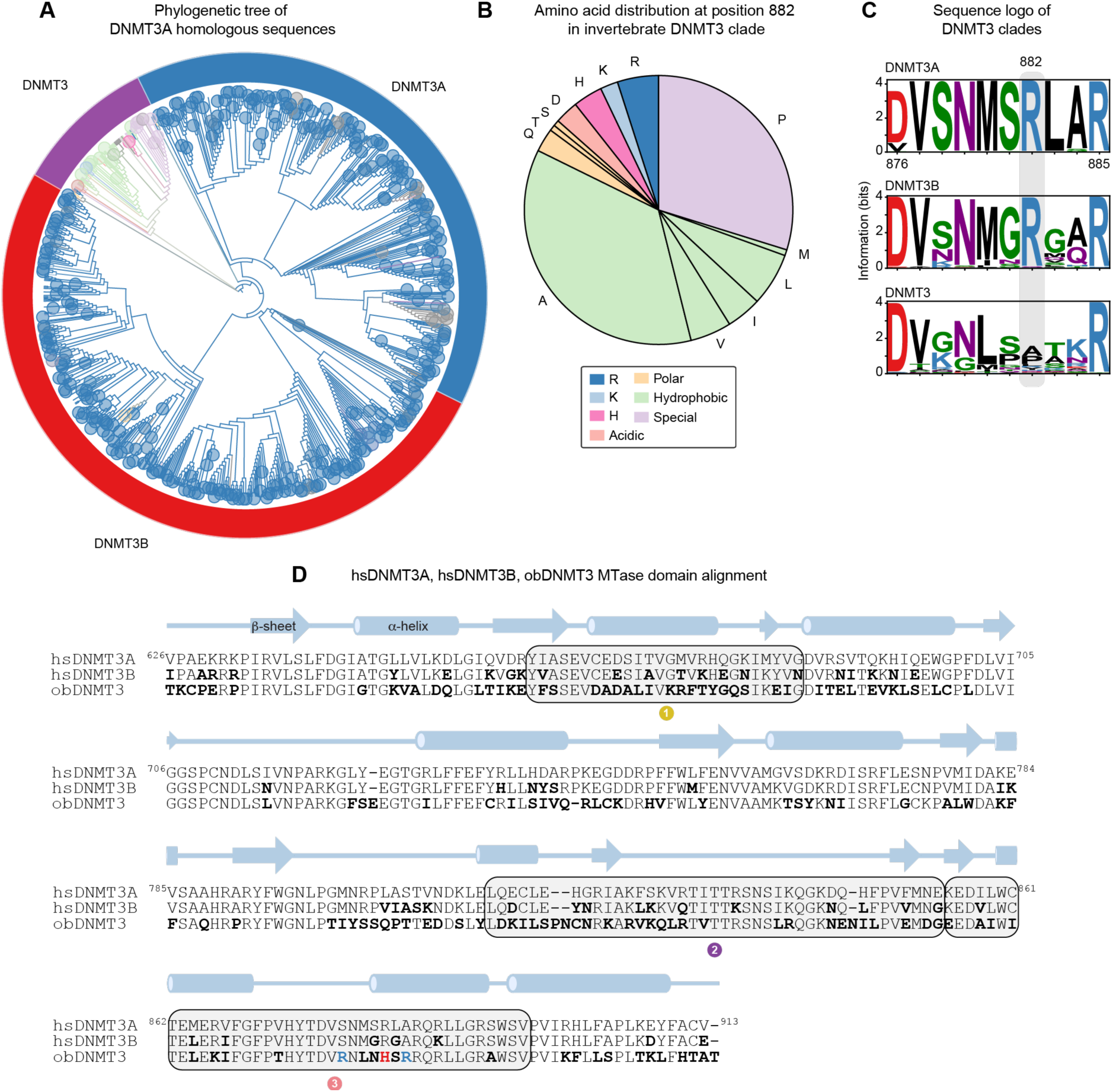
Phylogenetic analysis of DNMT3A. **J.** Cladogram of DNMT3A homologs in animals made from a maximum-likelihood phylogenetic tree generated with FastTree^48^ (v2.1.11) containing domains homologous to the PWWP, ADD, and MTase domain of hsDNMT3A (visualized with iToL v7.2.1). Branches and leaves are colored by position 882 chemistry. (color legend shared with panel **B**) **K.** Distribution of amino acids at the equivalent position to R882 in hsDNMT3A among invertebrate DNMT3s. **L.** Sequence logos of DNMT3A, DNMT3B, and invertebrate DNMT3 clades surrounding position R882. **M.** MTase sequence alignment of hsDNMT3A, hsDNMT3B, and obDNMT3. Bold indicates differences from hsDNMT3A. Grey boxes highlight regions perturbed in our deep mutational scan. Red highlights residue 882, blue highlights residues 878 and 884.

**Figure S2:**
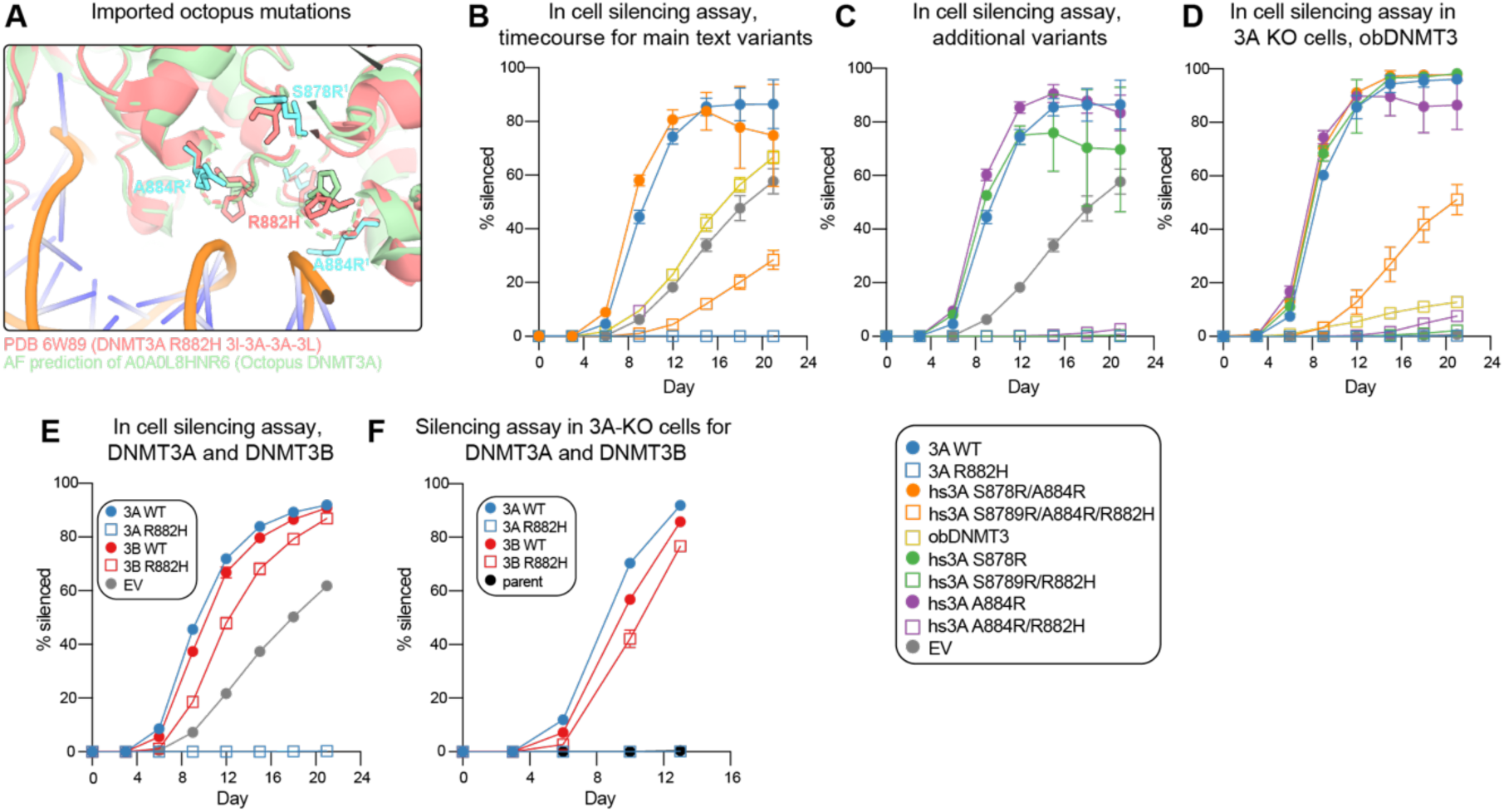
Additional in-cell silencing assays of phylogeny informed DNMT3A variants. **A.** Pymol overlay of hsDNMT3A^R882H^ (PDB 6w89) and an AlphaFold2 model ^54^ of obDNMT3. S878, R882, and A884 and their equivalent obDNMT3 residues are shown as sticks. **B.** In cell silencing assay of single and double octopus-informed mutants (complement to Figure 1D). **C.** In cell silencing assay of single and double octopus-informed mutants (additional variants associated with Figure 1D). **D.** In cell silencing assay of DNMT3B^WT^ and DNMT3B^R823H^ in DNMT3A knockout (KO) reporter cells. **E.** In cell silencing assay time course of DNMT3B^WT^ and DNMT3B^R823H^ (complement to Figure 1D) **F.** In cell silencing assay of DNMT3B^WT^ and DNMT3B^R823H^ in DNMT3A (KO) reporter cells. Data in **B–F** are mean ± S.D. with individual replicates shown (n=3).

**Figure S3:**
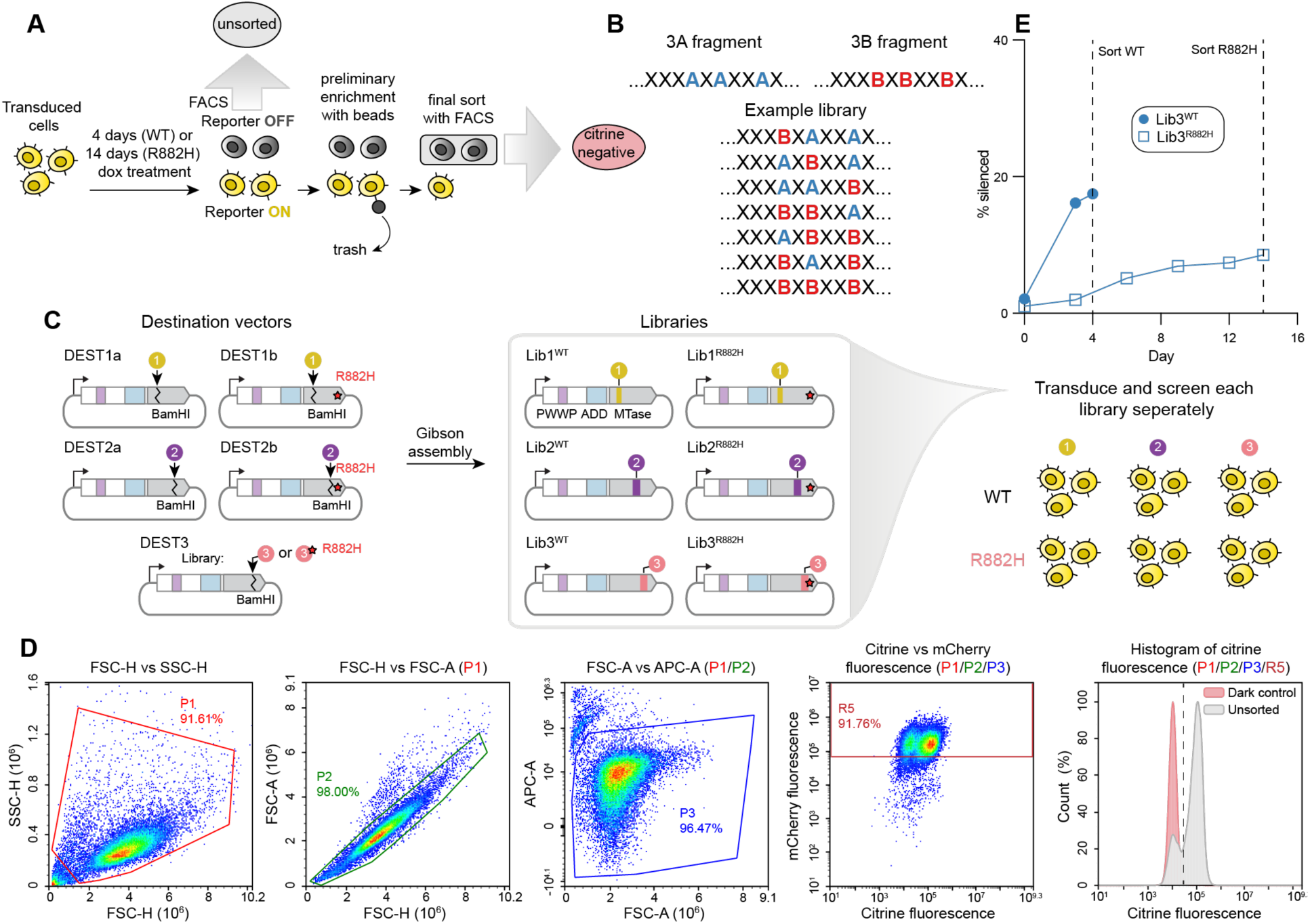
Additional deep mutational scanning methodology. **A.** Expanded sorting strategy for deep mutational scanning. Dashes represent surface marker^28^. Cells were silenced for 4 or 14 days. A portion of the initial pool of cells were sorted for mcherry positivity (to ensure proper reporter expression from the initial pool) and designated as “Unsorted”. The remaining pool of cells was then sorted first with a preliminary enrichment with magnetic beads and second by FACS. **B.** Schematic of DNMT3B informed library design. X=conserved residue between DNMT3A and DNMT3B, A=DNMT3A residue, B=DNMT3B residue. **C.** Cartoon representation of library cloning strategy. **D.** Representative flow cytometry gating strategy for silencing assays and DMS cell sorting. Helix NP NIR was used to sort viable cells. The citrine gate was set such that 1% of a control dark population of cells would be called as citrine+. **E.** Silencing of library 3 WT and library 3 R882H pools over time. Data are average of n=3 replicates.

**Figure S4:**
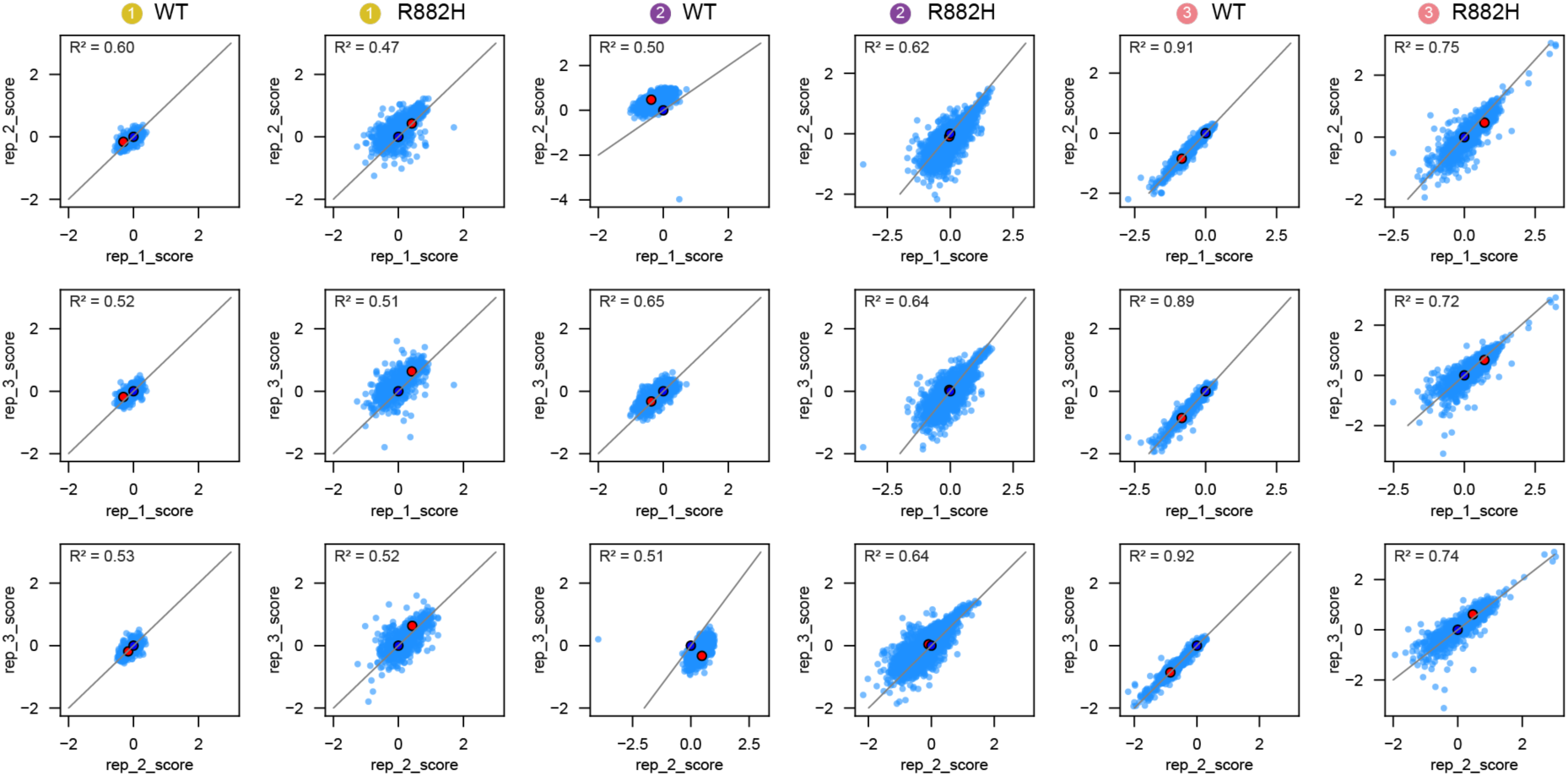
Library replicate correlation. Replicate correlation of each rep for each library. Blue dot=WT, Red dot= mean of nonsense mutations. Replicate 2 of library 2 WT was removed from further analysis due to poor correlation with replicates 1 and 3.

**Figure S5:**
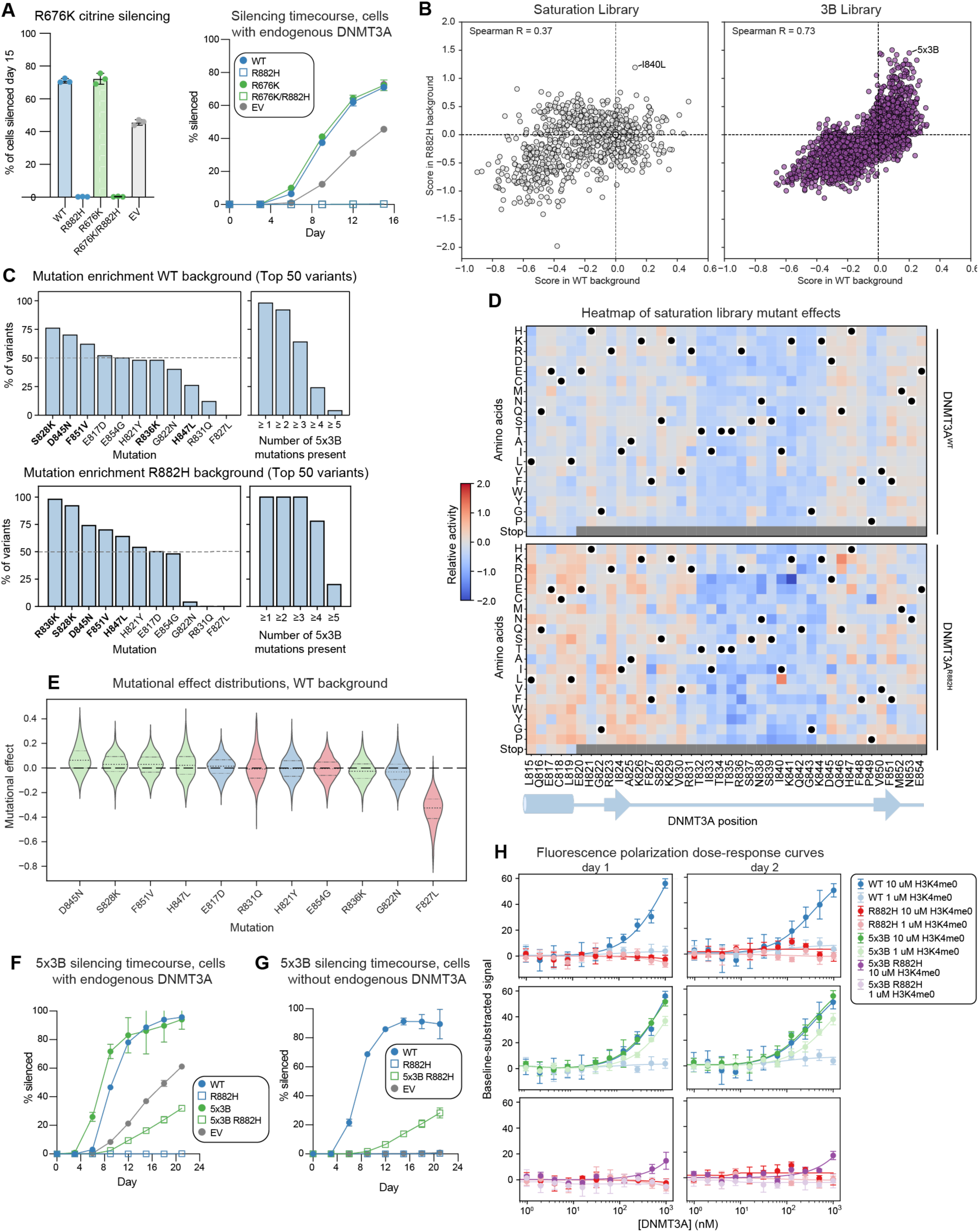
Additional analysis of TRD library (Library 2) **A.** In cell silencing assay of DNMT3A^R676K/R882H^ mutants after 15 days of dox treatment (left) or over time (right). **B.** Scatter plots of deep mutational scanning scores for DNMT3A^WT^ (*x* axis) or DNMT3A^R882H^ (*y* axis) for Library 2 (TRD) normalized to wild-type or R882H, respectively. Data split by saturation library (left) or 3B informed library (right). **C.** Mutational enrichment of individual DNMT3B informed mutations in the top 50 ranked variants in the WT background (top) or R882H background (bottom). Number of variants out of the top 50 scoring variants that contain the given mutation (left) and number of mutations in 5×3B in each of the top ranked variants (right). **D.** Heatmap of deep mutational scanning scores for the saturation library in the TRD library (library 2) for the WT (top) and R882H (bottom) backgrounds. **E.** Violin plot showing distribution of mutational effect for each of the DNMT3B informed mutations in the WT background. Mutational effect is calculated as the difference in score between matched variants with and without the named mutation (e.g. DNMT3A^variant X^ versus DNMT3A^variant X/R836K^). median and interquartile range are shown. Green= mutations in 5×3B, red= deleterious mutations, blue= other mutations. Mutations are numbered by DNMT3A residue. **F.** Time course of in cell silencing assay of 5×3B (complementary to Figure 2D). **G.** In cell silencing assay of DNMT3A^5x3B/R882H^ in DNMT3A KO reporter cells. **H.** Dose-response curves for fluorescence-polarization based DNA binding assay (Figure 2G). The experiment was performed twice in technical triplicate and both replicates are shown. Data in **B, C, D, and E** are mean of n = 3 biological replicates for R882H background libraries and mean of n=2 biological replicates for WT background libraries. The overall DMS experiment was performed once Data in **A, F, and G** are mean ± S.D. with individual replicates shown (n=3).

**Figure S6:**
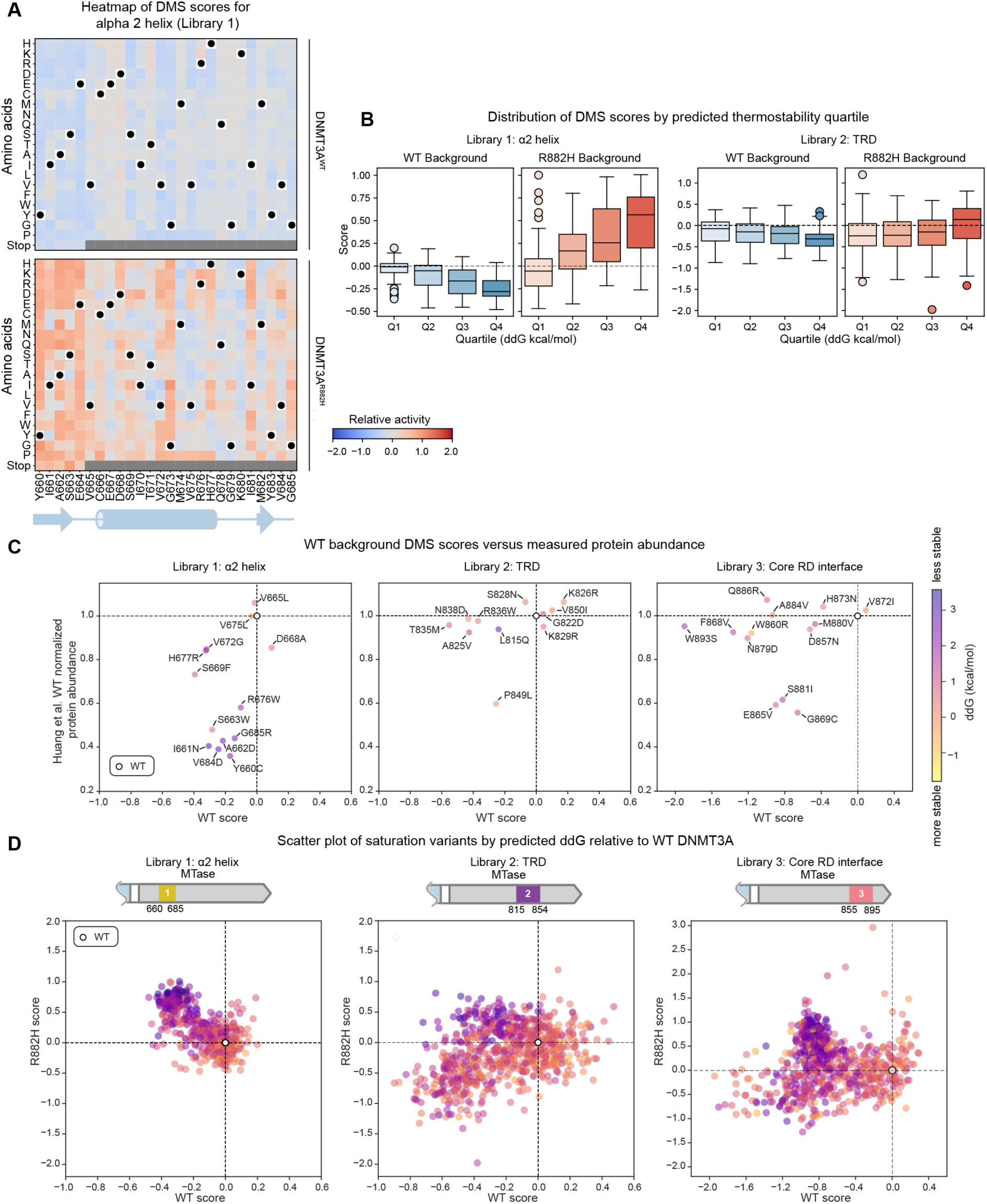
Comparison of deep mutational scanning data to predicted protein stability and measured protein abundance. **A.** Heatmap of deep mutational scanning scores for the saturation library in the WT (top) and R882H (bottom) backgrounds. **B.** Box plots of mutational scanning scores of each saturation library separated into quartiles based on ThermoMPNN^34^ predicted protein stability (Q1 most stable, Q4 least stable). ThermoMPNN predictions (**Table S2**) based on PDB 6w8b. **C.** Scatter plots of WT deep mutational scanning scores vs protein abundance measurements of clinical variants ^35^ colored by ThermoMPNN-predicted LlLlG of folding. Higher numbers indicate variants that are less stable than DNMT3A^WT^. **D.** Scatter plots of WT vs R882H background saturation library DMS data, colored by ThermoMPNN predicted LlLlG of folding compared to WT. DMS data in **A-D** are mean of n=3 biological replicates and the DMS was performed once.

**Figure S7:**
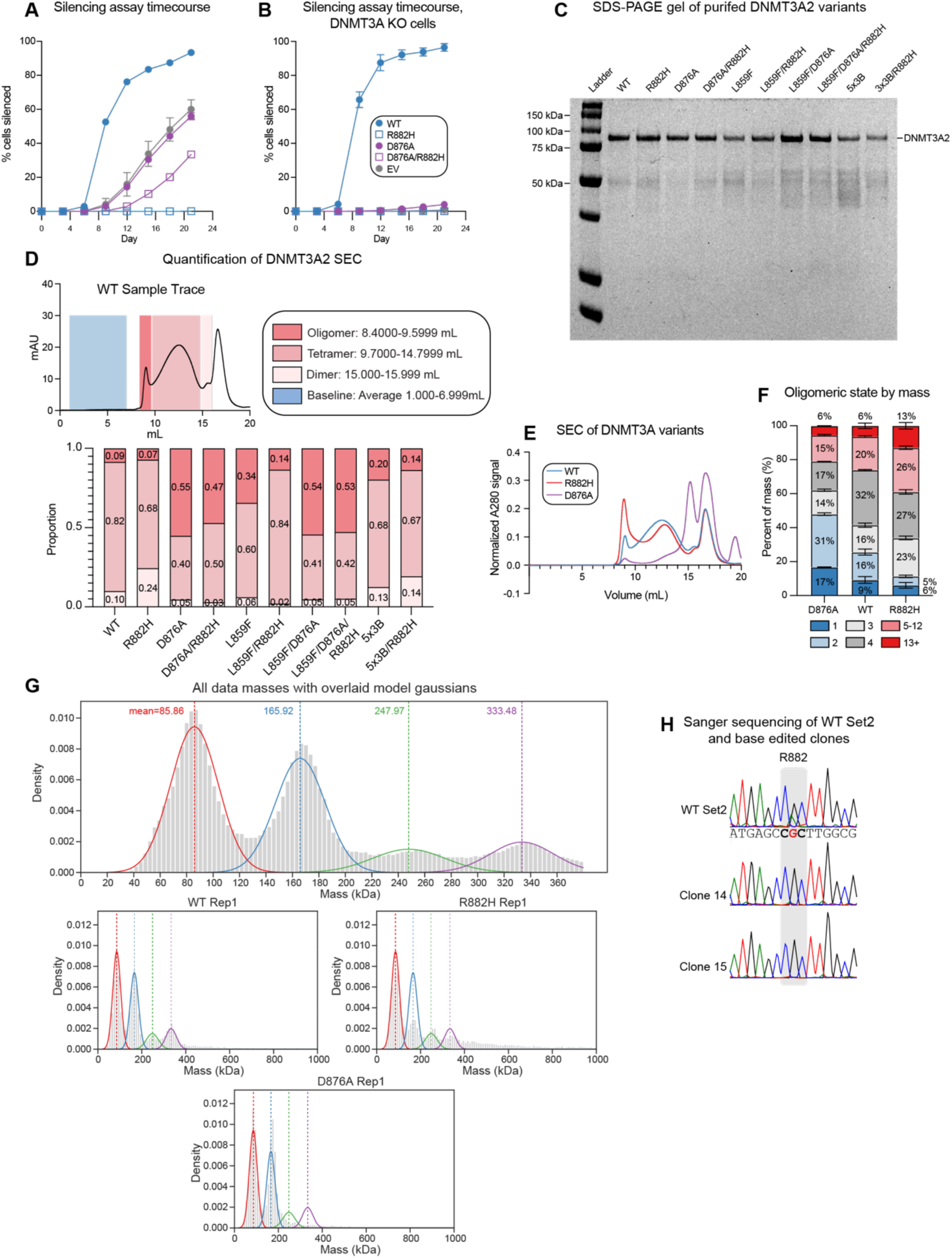
Additional biochemical validation of DNMT3A variants. **A.** Time course of in cell silencing assay of DNMT3A^D876A^, DNMT3A^D876A/R882H^, and controls (complementary to Figure 3D) **B.** In cell silencing assay of DNMT3A^D876A^ and DNMT3A^D876A/R882H^ in DNMT3A KO reporter cells. **C.** Coomassie stained SDS-PAGE gel of purified DNMT3A mutants (pre-SEC). **D.** Quantification of oligomeric state of DNMT3A variants by size exclusion chromatography (SEC) (Superose 6 10/300). Data are representative of n=2 replicates. **E.** Size exclusion chromatography (SEC) (Superose 6 10/300) traces of purified DNMT3A2 variants. Data are representative of n=2 technical replicates. **F.** Stacked bar plots showing the size distribution of pre-SEC purified DNMT3A proteins measured by mass photometry for indicated variants. n=3. **G.** (top) Histogram showing mass photometry of all replicates of all samples (3 replicates of each DNMT3A variant: WT, R882H, D876A) pooled and fit with a Gaussian mixture model (see Methods, Supplementary Code). Data same as in panel **F**. DNMT3A2 monomer actual mass= 80kDa. Model Gaussian means for each peak are shown. (bottom) Representative histogram of mass distribution for each variant with model gaussians calculated from pooled data overlaid. **H.** Sanger sequencing of the DNMT3A R882 locus of WT (top) and base edited (middle and bottom) Set 2 cells. Data in **A** and **B** are mean ± S.D. with individual replicates shown (n=3).

**Figure S8:**
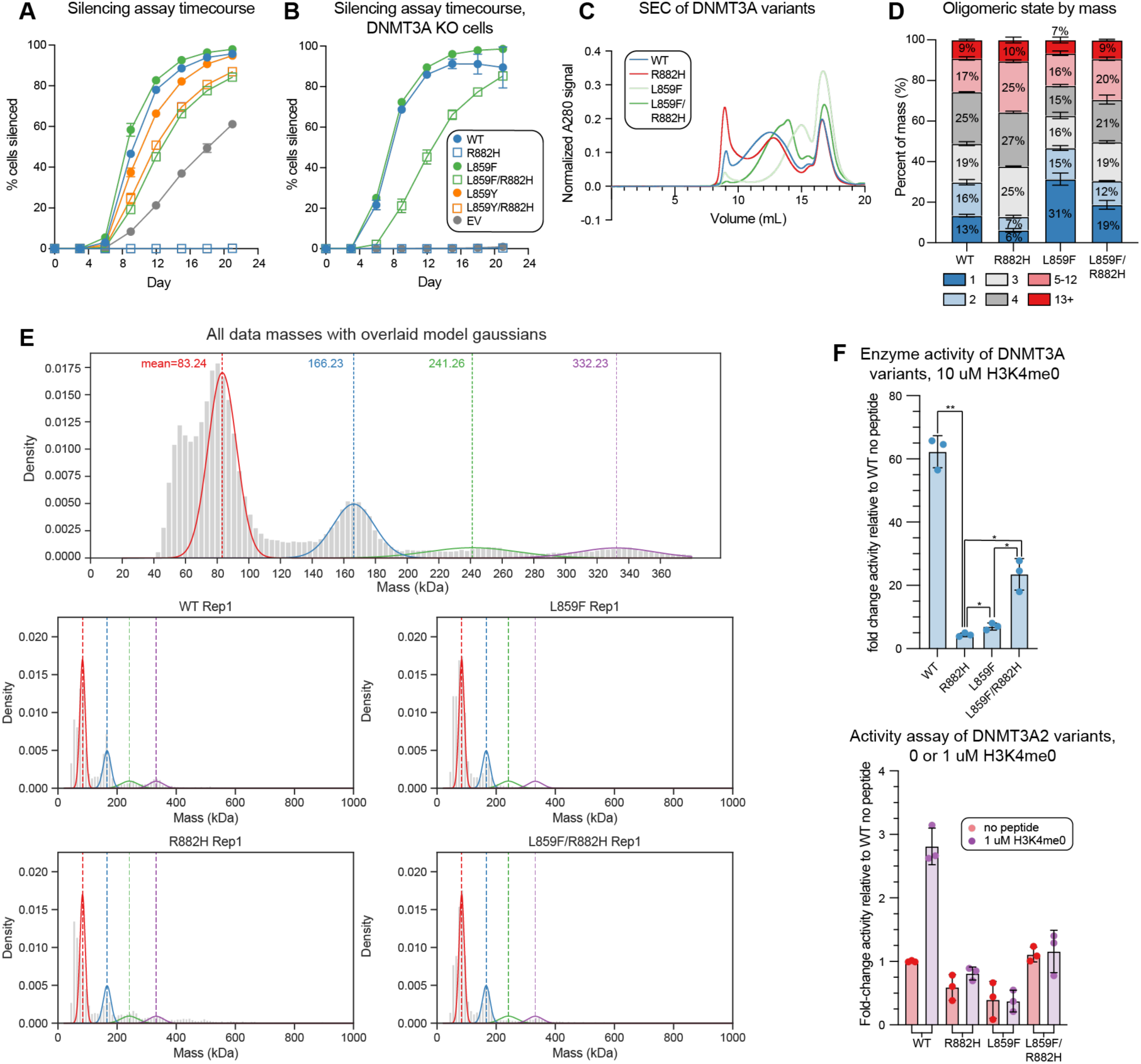
Additional biochemical validation of DNMT3A rescue variants. **A.** Time course of in cell silencing assay of DNMT3A^L859F^ and DNMT3A^L859Y^ and controls (complementary to Figure 4B) **B.** In cell silencing assay of DNMT3A^L859F^ and DNMT3A^L859F/R882H^ in DNMT3A KO reporter cells. **C.** Size exclusion chromatography (SEC) (Superose 6 10/300) traces of purified DNMT3A2 variants. Data are representative of n=2 technical replicates. **D.** Stacked bar plots showing the size distribution of pre-SEC purified DNMT3A proteins measured by mass photometry for indicated variants. n=3. **E.** (top) Histogram showing mass photometry of all replicates of all samples (3 replicates of each DNMT3A variant: WT, R882H, L859F, and L859F/R882H) pooled and fit with a Gaussian mixture model (see Methods, Supplementary Code). Data same as in panel D. DNMT3A2 monomer actual mass= 80kDa. Model Gaussian means for each peak are shown. (bottom) Representative histogram of mass distribution for each variant with model gaussians calculated from pooled data overlaid. **F.** Quantification of the activity of indicated DNMT3A variants for DNA methylation on a poly dI•dC substrate in the presence or absence of varying concentrations of H3K4me0. Data are mean ± S.D. with individual replicates shown (n=3) and are representative of two independent experiments. *P* values were calculated through two-tailed Welch’s *t* tests; * ≤ 0.05, ** ≤ 0.01, *** ≤ 0.001, *** ≤ 0.0001. Data in **A** and **B** are mean ± S.D. with individual replicates shown (n=3).

**Figure S9:**
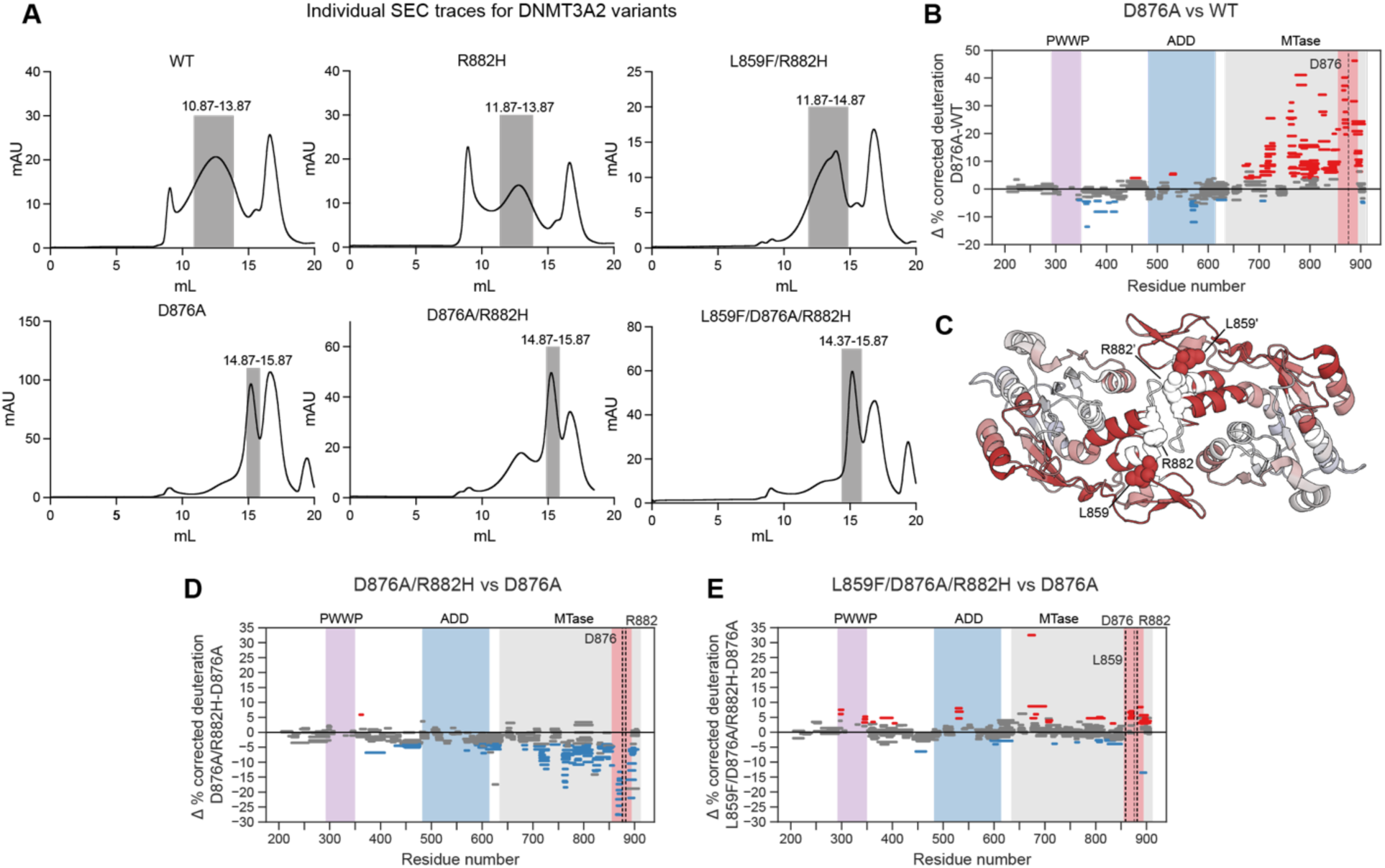
Additional data associated with HdX-MS. **A.** SEC traces of purified DNMT3A variants. Data are representative of n=2 technical replicates. Grey boxes indicate protein fractions pooled and concentrated for use in HdX-MS experiments. **B.** Woods plot showing differences in deuterium-labeled peptides (%) between DNMT3A^WT^ and DNMT3A^D876A^ at 100 sec. Significant differences in % deuterium uptake at the peptide level were identified using a dual-threshold approach: (1) a two-tailed Welch’s t-test at α = 0.05, and (2) a difference threshold of 2 times the pooled standard deviation from % deuterium uptake measurements across all peptides. Deprotected, protected, and non-significantly different peptides are shown in red, blue and grey respectively. **C.** Differential deuterium uptake between D876A and WT at 100 sec mapped on the inner MTase’ (PDB: 8tdr). Per-residue coloring was determined using the Keppel and Weis algorithm^38^. **D.** Woods plot showing deuterium differences (%) between D876A and D876A/R882H dimers at 100 sec. **E.** Woods plot showing deuterium differences (%) between DNMT3A^D876A^ and DNMT3A^L859F/D876A/R882H^ at 100 sec. Data in **B-E** are average of n = 4 technical replicates

**Figure S10:**
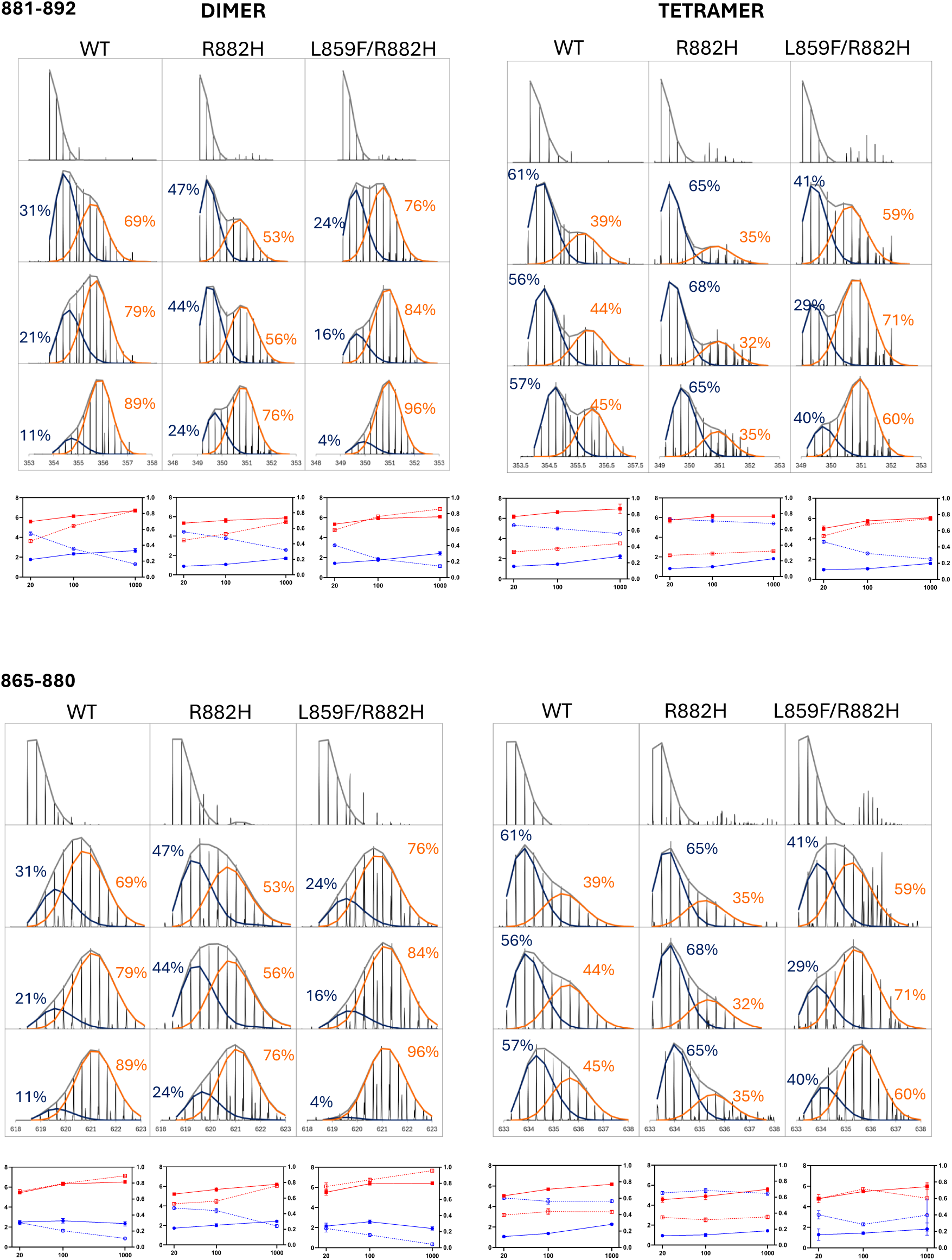
Bimodal deconvolution. Representative isotopic distributions and bimodal deconvolution across all time points (>99% confidence intervals) of selected peptides spanning the RD interface. Binomial fitting of the total population, the ‘closed’ state and the ‘open’ state is shown in grey, dark blue and orange respectively, along with the % fraction of each population. Line plots correspond to the D-uptake of the ‘closed’ (blue) and ‘open’ (red) states (n=4, left y-axis). Dashed lines correspond to the % fraction of each population (right y-axis).

## Acknowledgements

We thank current and former members of the Liau Lab for helpful discussions and comments on the manuscript, in particular A. Waterbury, H. S. Kwok, C. Lee, O. Zhang, J. W. Morriss, and S. P. Shen. We thank J. Nelson and C. Maesner for assistance with FACS, K. Arnett and the Center for Macromolecular Interactions at Harvard Medical School for training and assistance with Mass Photometry, D. Bolduc for biochemistry advice, and S. P. Berry for helpful discussions and comments on the manuscript as well as computational advice. We thank members of the Proteomics Group at the Broad Institute of MIT & Harvard for insightful discussions and our gratitude to Vlad Sarpe at Trajan Scientific and Medical for software support.

## Author Contributions

E.M.G. and B.B.L. conceived the study and wrote the manuscript with assistance from O.R.L. and M.P.. E.M.G., O.R.L., S.R., N.Z.L., M.P. and B.B.L. designed experiments and analyzed data. E.M.G. performed deep mutational scanning and analysis, cell biology experiments, gene editing experiments, biochemistry experiments, and phylogenetic analysis. O.R.L performed cell culture and biochemistry experiments. S.R. designed, performed, and analyzed HDX-MS experiments with assistance from M.M. and input from M.P.. N.Z.L. performed biochemistry experiments. J.K.L. performed phylogenetic analysis and cell biology experiments. D.D.H. performed biochemistry experiments. M.U.Z. and L.W.C. assisted in gene editing experiments. M.P. oversaw HDX-MS experiments with input from S.A.C.. B.B.L. supervised and held overall responsibility for the study.

## Declaration of Interests

S.A.C. is a member of the scientific advisory boards of Kymera, Stand Up to Cancer, PTM BioLabs and PrognomIQ. B.B.L. is a cofounder, shareholder, and member of the scientific advisory board of Light Horse Therapeutics and receives research funding from Ono Pharmaceutical Co., Ltd.. The remaining authors declare no competing interests.

N.Z.L. is currently affiliated with the California Institute for Quantitative Biosciences (QB3) at the University of California, Berkeley. J.K.L. is currently affiliated with the Biophysics Graduate Program at Stanford University.

## Data availability

The original mass spectra and the protein sequence databases used for searches have been deposited in the public proteomics repository MassIVE (http://massive.ucsd.edu) and will be made public upon acceptance of the manuscript. Sequencing data from the DNMT3A and DNMT3A^R882H^ deep mutational scans have been deposited in the NCBI Sequence Read Archive (SRA) and will be made public upon acceptance of the manuscript. Code associated with this study can be found at https://github.com/liaulab/DNMT3A_2025.

## References

1. Garcia-Seisdedos, H., Empereur-Mot, C., Elad, N., and Levy, E.D. (2017). Proteins evolve on the edge of supramolecular self-assembly. Nature 548, 244–247. 10.1038/nature23320.

2. Jaiswal, S., Fontanillas, P., Flannick, J., Manning, A., Grauman, P.V., Mar, B.G., Lindsley, R.C., Mermel, C.H., Burtt, N., Chavez, A., et al. (2014). Age-Related Clonal Hematopoiesis Associated with Adverse Outcomes. New Engl. J. Med. 371, 2488–2498. 10.1056/nejmoa1408617.

3. Genovese, G., K., K.A., E., H.R., Johan, L., A., R.S., F., B.S., Kimberly, C., Eran, M., M., N.B., Menachem, F., et al. (2014). Clonal Hematopoiesis and Blood-Cancer Risk Inferred from Blood DNA Sequence. New Engl. J. Med. 371, 2477–2487. 10.1056/nejmoa1409405.

4. Brunetti, L., Gundry, M.C., and Goodell, M.A. (2017). DNMT3A in Leukemia. Cold Spring Harb Perspect Med 7, a030320. 10.1101/cshperspect.a030320.

5. Russler-Germain, D.A., Spencer, D.H., Young, M.A., Lamprecht, T.L., Miller, C.A., Fulton, R., Meyer, M.R., Erdmann-Gilmore, P., Townsend, R.R., Wilson, R.K., et al. (2014). The R882H DNMT3A mutation associated with AML dominantly inhibits wild-type DNMT3A by blocking its ability to form active tetramers. Cancer Cell 25, 442–454. 10.1016/j.ccr.2014.02.010.

6. Kim, S.J., Zhao, H., Hardikar, S., Singh, A.K., Goodell, M.A., and Chen, T. (2013). A DNMT3A mutation common in AML exhibits dominant-negative effects in murine ES cells. Blood 122, 4086–4089. 10.1182/blood-2013-02-483487.

7. Challen, G.A., and Goodell, M.A. (2020). Clonal hematopoiesis: mechanisms driving dominance of stem cell clones. Blood 136, 1590–1598. 10.1182/blood.2020006510.

8. Lu, J., Guo, Y., Yin, J., Chen, J., Wang, Y., Wang, G.G., and Song, J. (2024). Structure-guided functional suppression of AML-associated DNMT3A hotspot mutations. Nat Commun 15, 3111. 10.1038/s41467-024-47398-y.

9. Anteneh, H., Fang, J., and Song, J. (2020). Structural basis for impairment of DNA methylation by the DNMT3A R882H mutation. Nat Commun 11, 2294. 10.1038/s41467-020-16213-9.

10. Holz-Schietinger, C., Matje, D.M., and Reich, N.O. (2012). Mutations in DNA Methyltransferase (DNMT3A) Observed in Acute Myeloid Leukemia Patients Disrupt Processive Methylation. J Biol Chem 287, 30941–30951. 10.1074/jbc.m112.366625.

11. Nguyen, T.-V., Yao, S., Wang, Y., Rolfe, A., Selvaraj, A., Darman, R., Ke, J., Warmuth, M., Smith, P.G., Larsen, N.A., et al. (2019). The R882H DNMT3A hot spot mutation stabilizes the formation of large DNMT3A oligomers with low DNA methyltransferase activity. J Biol Chem 294, 16966–16977. 10.1074/jbc.ra119.010126.

12. Mack, A., Emperle, M., Schnee, P., Adam, S., Pleiss, J., Bashtrykov, P., and Jeltsch, A. (2022). Preferential Self-interaction of DNA Methyltransferase DNMT3A Subunits Containing the R882H Cancer Mutation Leads to Dominant Changes of Flanking Sequence Preferences. J. Mol. Biol. 434, 167482. 10.1016/j.jmb.2022.167482.

13. Emperle, M., Rajavelu, A., Kunert, S., Arimondo, P.B., Reinhardt, R., Jurkowska, R.Z., and Jeltsch, A. (2018). The DNMT3A R882H mutant displays altered flanking sequence preferences. Nucleic Acids Res 46, 3130–3139. 10.1093/nar/gky168.

14. Garcia, E.M., Lue, N.Z., Liang, J.K., Lieberman, W.K., Hwang, D.D., Woods, J.C., and Liau, B.B. (2023). Base Editor Scanning Reveals Activating Mutations of DNMT3A. ACS Chem. Biol. 18, 2030–2038. 10.1021/acschembio.3c00257.

15. Lu, R., Wang, J., Ren, Z., Yin, J., Wang, Y., Cai, L., and Wang, G.G. (2019). A Model System for Studying the DNMT3A Hotspot Mutation (DNMT3AR882) Demonstrates a Causal Relationship between Its Dominant-Negative Effect and Leukemogenesis. Cancer Res 79, 3583–3594. 10.1158/0008-5472.can-18-3275.

16. Emperle, M., Adam, S., Kunert, S., Dukatz, M., Baude, A., Plass, C., Rathert, P., Bashtrykov, P., and Jeltsch, A. (2019). Mutations of R882 change flanking sequence preferences of the DNA methyltransferase DNMT3A and cellular methylation patterns. Nucleic Acids Research 47, 11355–11367. 10.1093/nar/gkz911.

17. Norvil, A.B., AlAbdi, L., Liu, B., Tu, Y.H., Forstoffer, N.E., Michie, A.R., Chen, T., and Gowher, H. (2020). The acute myeloid leukemia variant DNMT3A Arg882His is a DNMT3B-like enzyme. Nucleic Acids Research 48, 3761–3775. 10.1093/nar/gkaa139.

18. Emperle, M., Dukatz, M., Kunert, S., Holzer, K., Rajavelu, A., Jurkowska, R.Z., and Jeltsch, A. (2018). The DNMT3A R882H mutation does not cause dominant negative effects in purified mixed DNMT3A/R882H complexes. Sci Rep 8, 13242. 10.1038/s41598-018-31635-8.

19. Jia, D., Jurkowska, R.Z., Zhang, X., Jeltsch, A., and Cheng, X. (2007). Structure of Dnmt3a bound to Dnmt3L suggests a model for de novo DNA methylation. Nature 449, 248–251. 10.1038/nature06146.

20. Lin, C.-C., Chen, Y.-P., Yang, W.-Z., Shen, J.C.K., and Yuan, H.S. (2020). Structural insights into CpG-specific DNA methylation by human DNA methyltransferase 3B. Nucleic Acids Res. 48, 3949–3961. 10.1093/nar/gkaa111.

21. Zhang, Z.-M., Lu, R., Wang, P., Yu, Y., Chen, D., Gao, L., Liu, S., Ji, D., Rothbart, S.B., Wang, Y., et al. (2018). Structural basis for DNMT3A-mediated de novo DNA methylation. Nature 554, 387–391. 10.1038/nature25477.

22. Watson, C.J., Papula, A.L., Poon, G.Y.P., Wong, W.H., Young, A.L., Druley, T.E., Fisher, D.S., and Blundell, J.R. (2020). The evolutionary dynamics and fitness landscape of clonal hematopoiesis. Science 367, 1449–1454. 10.1126/science.aay9333.

23. J., L.T., Li, D., J., W.M., D., M.M., Tamara, L., E., L.D., Cyriac, K., E., P.J., Jack, B., John, W., et al. (2010). DNMT3A Mutations in Acute Myeloid Leukemia. N. Engl. J. Med. 363, 2424–2433. 10.1056/nejmoa1005143.

24. Garcia-Seisdedos, H., Villegas, J.A., and Levy, E.D. (2019). Infinite Assembly of Folded Proteins in Evolution, Disease, and Engineering. Angew. Chem. Int. Ed. 58, 5514–5531. 10.1002/anie.201806092.

25. Bintu, L., Yong, J., Antebi, Y.E., McCue, K., Kazuki, Y., Uno, N., Oshimura, M., and Elowitz, M.B. (2016). Dynamics of epigenetic regulation at the single-cell level. Science 351, 720–724. 10.1126/science.aab2956.

26. Lue, N.Z., Garcia, E.M., Ngan, K.C., Lee, C., Doench, J.G., and Liau, B.B. (2022). Base editor scanning charts the DNMT3A activity landscape. Nat. Chem. Biol. 19, 176–186. 10.1038/s41589-022-01167-4.

27. Norvil, A.B., Petell, C.J., Alabdi, L., Wu, L., Rossie, S., and Gowher, H. (2018). Dnmt3b Methylates DNA by a Noncooperative Mechanism, and Its Activity Is Unaffected by Manipulations at the Predicted Dimer Interface. Biochemistry 57, 4312–4324. 10.1021/acs.biochem.6b00964.

28. Tycko, J., DelRosso, N., Hess, G.T., Aradhana, Banerjee, A., Mukund, A., Van, M.V., Ego, B.K., Yao, D., Spees, K., et al. (2020). High-Throughput Discovery and Characterization of Human Transcriptional Effectors. Cell 183, 2020–2035.e16. 10.1016/j.cell.2020.11.024.

29. Rubin, A.F., Gelman, H., Lucas, N., Bajjalieh, S.M., Papenfuss, A.T., Speed, T.P., and Fowler, D.M. (2017). A statistical framework for analyzing deep mutational scanning data. Genome Biol. 18, 150. 10.1186/s13059-017-1272-5.

30. Gao, L., Emperle, M., Guo, Y., Grimm, S.A., Ren, W., Adam, S., Uryu, H., Zhang, Z.-M., Chen, D., Yin, J., et al. (2020). Comprehensive structure-function characterization of DNMT3B and DNMT3A reveals distinctive de novo DNA methylation mechanisms. Nat. Commun. 11, 3355. 10.1038/s41467-020-17109-4.

31. Dukatz, M., Dittrich, M., Stahl, E., Adam, S., Mendoza, A. de, Bashtrykov, P., and Jeltsch, A. (2022). DNA methyltransferase DNMT3A forms interaction networks with the CpG site and flanking sequence elements for efficient methylation. J. Biol. Chem. 298, 102462. 10.1016/j.jbc.2022.102462.

32. Mallona, I., Ilie, I.M., Karemaker, I.D., Butz, S., Manzo, M., Caflisch, A., and Baubec, T. (2020). Flanking sequence preference modulates de novo DNA methylation in the mouse genome. Nucleic Acids Res 49, 145–157. 10.1093/nar/gkaa1168.

33. Guo, X., Wang, L., Li, J., Ding, Z., Xiao, J., Yin, X., He, S., Shi, P., Dong, L., Li, G., et al. (2015). Structural insight into autoinhibition and histone H3-induced activation of DNMT3A. Nature 517, 640–644. 10.1038/nature13899.

34. Dieckhaus, H., Brocidiacono, M., Randolph, N.Z., and Kuhlman, B. (2024). Transfer learning to leverage larger datasets for improved prediction of protein stability changes. Proc. Natl. Acad. Sci. United States Am. 121, e2314853121. 10.1073/pnas.2314853121.

35. Huang, Y.-H., Chen, C.-W., Sundaramurthy, V., Słabicki, M., Hao, D., Watson, C.J., Tovy, A., Reyes, J.M., Dakhova, O., Crovetti, B.R., et al. (2022). Systematic Profiling of DNMT3A Variants Reveals Protein Instability Mediated by the DCAF8 E3 Ubiquitin Ligase Adaptor. Cancer Discovery 12, 220–235. 10.1158/2159-8290.cd-21-0560.

36. Tristan, C., Shahani, N., Sedlak, T.W., and Sawa, A. (2011). The diverse functions of GAPDH: Views from different subcellular compartments. Cell. Signal. 23, 317–323. 10.1016/j.cellsig.2010.08.003.

37. Quentmeier, H., Pommerenke, C., Dirks, W.G., Fähnrich, S., Hauer, V., Uphoff, C.C., Zaborski, M., and Drexler, H.G. (2020). DNMT3A R882H mutation in acute myeloid leukemia cell line SET-2. Leuk. Res. 88, 106270. 10.1016/j.leukres.2019.106270.

38. Keppel, T.R., and Weis, D.D. (2015). Mapping Residual Structure in Intrinsically Disordered Proteins at Residue Resolution Using Millisecond Hydrogen/Deuterium Exchange and Residue Averaging. J. Am. Soc. Mass Spectrom. 26, 547–554. 10.1007/s13361-014-1033-6.

39. Livesey, B.J., and Marsh, J.A. (2022). The properties of human disease mutations at protein interfaces. PLoS Comput. Biol. 18, e1009858. 10.1371/journal.pcbi.1009858.

40. Fu, H., Mo, X., and Ivanov, A.A. (2025). Decoding the functional impact of the cancer genome through protein–protein interactions. Nat. Rev. Cancer 25, 189–208. 10.1038/s41568-024-00784-6.

41. Schweke, H., Pacesa, M., Levin, T., Goverde, C.A., Kumar, P., Duhoo, Y., Dornfeld, L.J., Dubreuil, B., Georgeon, S., Ovchinnikov, S., et al. (2024). An atlas of protein homo-oligomerization across domains of life. Cell 187, 999–1010.e15. 10.1016/j.cell.2024.01.022.

42. Xie, X., Zhang, O., Yeo, M.J.R., Lee, C., Tao, R., Harry, S.A., Payne, N.C., Nam, E., Paul, L., Li, Y., et al. (2025). Converging mechanism of UM171 and KBTBD4 neomorphic cancer mutations. Nature 639, 241–249. 10.1038/s41586-024-08533-3.

43. Seisdedos, H.G., Levin, T., Shapira, G., Freud, S., and Levy, E.D. (2022). Mutant libraries reveal negative design shielding proteins from supramolecular self-assembly and relocalization in cells. Proc. Natl. Acad. Sci. 119, e2101117119. 10.1073/pnas.2101117119.

44. Martin, M. (2011). Cutadapt removes adapter sequences from high-throughput sequencing reads | Martin | EMBnet. journal. EMBnet.journal 17, 10–12. 10.14806/ej.17.1.200.

45. Bushnell, B., Rood, J., and Singer, E. (2017). BBMerge – Accurate paired shotgun read merging via overlap. PLoS ONE 12, e0185056. 10.1371/journal.pone.0185056.

46. Eddy, S.R. (2011). Accelerated Profile HMM Searches. PLoS Comput. Biol. 7, e1002195. 10.1371/journal.pcbi.1002195.

47. Sievers, F., Wilm, A., Dineen, D., Gibson, T.J., Karplus, K., Li, W., Lopez, R., McWilliam, H., Remmert, M., Söding, J., et al. (2011). Fast, scalable generation of high-quality protein multiple sequence alignments using Clustal Omega. Mol. Syst. Biol. 7, MSB201175. 10.1038/msb.2011.75.

48. Price, M.N., Dehal, P.S., and Arkin, A.P. (2009). FastTree: Computing Large Minimum Evolution Trees with Profiles instead of a Distance Matrix. Mol. Biol. Evol. 26, 1641–1650. 10.1093/molbev/msp077.

49. Berry, S.P., and Ganesh, S. (2023). bioviper.

50. Tareen, A., and Kinney, J.B. (2020). Logomaker: beautiful sequence logos in Python. Bioinformatics 36, 2272–2274. 10.1093/bioinformatics/btz921.

51. Hummels, K.R., Berry, S.P., Li, Z., Taguchi, A., Min, J.K., Walker, S., Marks, D.S., and Bernhardt, T.G. (2023). Coordination of bacterial cell wall and outer membrane biosynthesis. Nature 615, 300–304. 10.1038/s41586-023-05750-0.

52. Huerta-Cepas, J., Serra, F., and Bork, P. (2016). ETE 3: Reconstruction, Analysis, and Visualization of Phylogenomic Data. Mol. Biol. Evol. 33, 1635–1638. 10.1093/molbev/msw046.

53. Letunic, I., and Bork, P. (2024). Interactive Tree of Life (iTOL) v6: recent updates to the phylogenetic tree display and annotation tool. Nucleic Acids Res. 52, W78–W82. 10.1093/nar/gkae268.

54. Jumper, J., Evans, R., Pritzel, A., Green, T., Figurnov, M., Ronneberger, O., Tunyasuvunakool, K., Bates, R., Žídek, A., Potapenko, A., et al. (2021). Highly accurate protein structure prediction with AlphaFold. Nature 596, 583–589. 10.1038/s41586-021-03819-2.

